# Macrophages transfer mitochondria to sensory neurons to resolve inflammatory pain

**DOI:** 10.1101/2020.02.12.940445

**Authors:** Michiel van der Vlist, Ramin Raoof, Hanneke L.D.M. Willemen, Judith Prado, Sabine Versteeg, Christian Martin Gil, Martijn Vos, Roeland E. Lokhorst, R. Jeroen Pasterkamp, Toshiyuki Kojima, Hajime Karasuyama, William Khoury-Hanold, Linde Meyaard, Niels Eijkelkamp

## Abstract

The current paradigm is that inflammatory pain passively resolves following the cessation of inflammation. Yet, in a substantial proportion of patients with inflammatory diseases, resolution of inflammation is not sufficient to resolve pain, resulting in chronic pain. Mechanistic insight how inflammatory pain is resolved is lacking. Here we show that macrophages actively control resolution of inflammatory pain remotely from the site of inflammation by transferring mitochondria to sensory neurons. During resolution of inflammatory pain in mice, M2-like macrophages infiltrate the dorsal root ganglia that contain the somata of sensory neurons, concurrent with the recovery of oxidative phosphorylation in sensory neurons. The resolution of pain and the transfer of mitochondria requires expression of CD200 Receptor (CD200R) on macrophages and the non-canonical CD200R-ligand iSec1 on sensory neurons. Our data reveal a novel mechanism for active resolution of inflammatory pain.

**Graphical Abstract:** 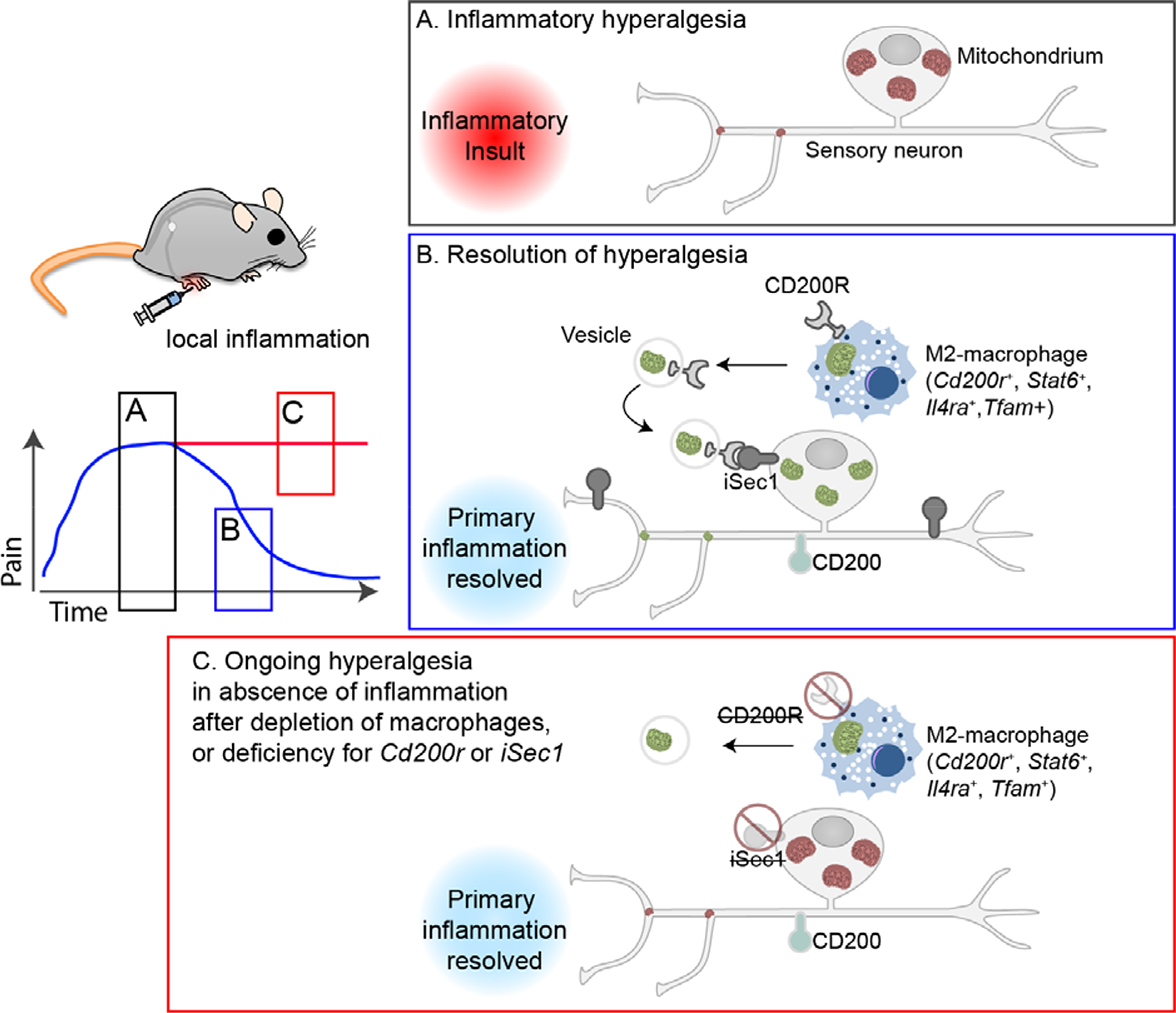

## Introduction

Pain and pain hypersensitivity (hyperalgesia) are functional features of inflammation that serve to protect the tissue from further damage. At the site of inflammation, immune cells and inflammatory mediators, such as IL-1β, TNF, and bradykinin, sensitize and activate sensory neurons, which cause pain and hyperalgesia (Ghasemlou et al., 2015; Peng et al., 2016). While the initiation of inflammatory pain is relatively well understood (Basbaum et al., 2009; Ji et al., 2016), the mechanisms of inflammatory pain resolution are less well characterized. Resolution of inflammatory pain is often considered to be the direct result of waning of inflammation. However, in a substantial proportion of patients with inflammatory diseases, such as rheumatoid arthritis and inflammatory bowel disease, spontaneous or treatment-induced resolution of inflammation does not reduce pain (Bielefeldt et al.; Hughes et al.; Krock et al., 2018; Lee et al., 2011; Lomholt et al.). Basic discovery research to understand mechanisms of endogenous pain-resolution may help us understand how chronic pain develops when resolution pathways fail (Price et al., 2018).

Macrophages are immune cells with enormous plasticity and are well known for their ability to induce tissue healing and resolution of inflammation (Wynn and Vannella, 2016). Macrophages are strongly imprinted by their tissue of residence (Gautier et al., 2012; Lavin et al., 2014). Peripheral nervous tissue shapes resident macrophages to have unique features compared to microglia and/or macrophages outside the nervous system (Kolter et al., 2020; Wang et al., 2020; Ydens et al., 2020). After nerve damage, monocyte-derived macrophages engraft nervous tissue (Ydens et al., 2020), are skewed by sensory neurons into an M1-like phenotype (Simeoli et al., 2017), and accumulate in the DRG to initiate and maintain neuropathic pain (Yu et al., 2020). Thus, nervous tissue macrophages contribute to neuropathic pain. However, because macrophages can contribute to tissue healing and resolution of inflammation as well as neuropathic pain (Niehaus et al., 2021), we here set out to better understand the endogenous mechanisms for resolution of inflammatory pain and the role of macrophages in this process, using transient inflammatory pain models.

## Results

We injected λ−carrageenan into the hind paw of mice (intraplantar; i.pl.) as a model for transient inflammatory pain (Supplemental fig. 1A) (Winter and Flataker, 1965). Treated mice displayed signs of pain hypersensitivity, such as allodynia/hyperalgesia as assessed by the von Frey and Hargreaves tests, and postural changes measured with dynamic weight bearing. Carrageenan-induced hyperalgesia resolved within ∼3-4 days (Fig. 1A). We analysed the cellular composition of lumbar (L3-L5) dorsal root ganglia (DRG) which contain the somata of sensory neurons innervating the hind paw and observed an accumulation of macrophages. Macrophage numbers peaked at day 3 and returned to baseline levels after resolution of inflammatory hyperalgesia (Fig. 1B/C, supplemental figs. 1B, and supplemental movies 1 and 2). Infiltration of macrophages was specific to the DRG that innervate the inflamed paw, and was not observed at the contralateral side (supplemental fig. 1C). In contrast, during the entire course of inflammatory hyperalgesia, T cells, B cells or other CD45^+^ immune cell numbers in the DRG did not change significantly (Figs. 1B and supplemental fig. 1B). To address the function of these macrophages in pain resolution, we selectively depleted monocytes and macrophages by intraperitoneal (i.p.) injection of diphtheria toxin (DT) in *Lysm*^cre/+^ x *Csf1r*^DTR/+^ mice (Schreiber et al., 2013) (from here on referred to as ‘MM^dtr^’). MM^dtr^ mice were compared with *Lysm^+/+^Csf1r^LsL-DTR/+^* littermates as control (from here on “Ctr”). DT treatment was started one day prior to i.pl. injection of carrageenan, and repeated daily. DT administration depleted monocytes and macrophages in the DRG, spinal cord, paw and blood (Supplemental figs. 2A-E) but did not affect the number and morphology of microglia in spinal cord (Supplemental figs. 2G-J). The induction and magnitude of carrageenan-induced hyperalgesia and inflammation in these mice was comparable to littermate controls (Fig. 1D and supplemental fig. 2K). However, MM^dtr^ mice failed to resolve inflammatory mechanical hyperalgesia (Fig. 1D), thermal hyperalgesia (Supplemental fig. 3A) and postural changes related to inflammatory pain for at least six days (Fig. 1E) in both male and female mice (Fig. 1F). Similarly, MM^dtr^ mice failed to resolve Complete Freund’s Adjuvant (CFA)-induced transient inflammatory hyperalgesia for at least 12 days (Fig. 1G, and supplemental fig. 3B).

**Figure 1.**
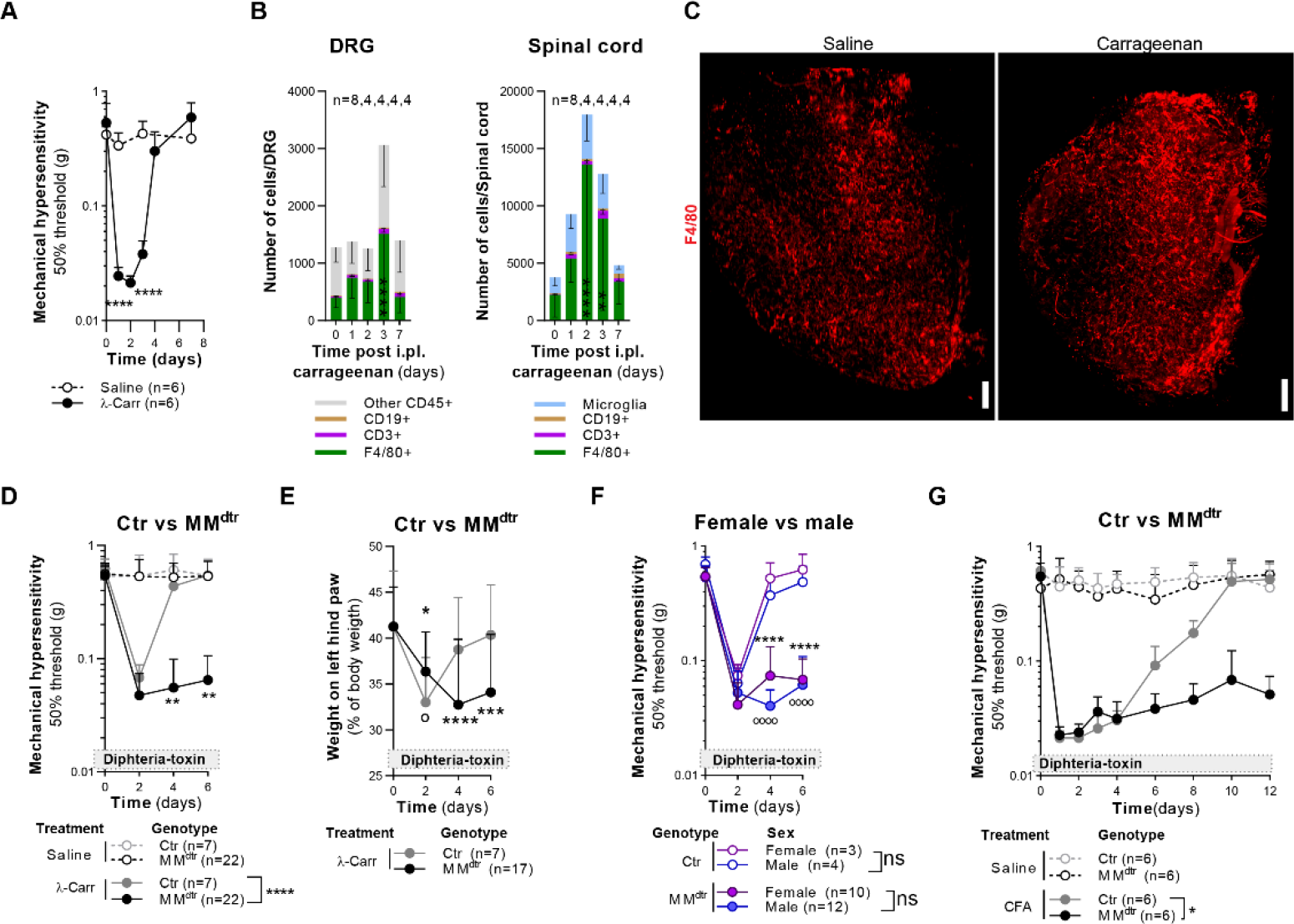
Monocytes/macrophages are required to resolve inflammatory pain. (A) Course of mechanical hyperalgesia after i.pl. injection of 1% carrageenan in the left hind paw and saline in the right hind paw. Statistics tested by multiple t test. (B) Absolute number of CD45^+^ cells classified to subset per lumbar dorsal root ganglia (DRG, L3-L5) and spinal cord of mice that received 1% carrageenan. See Supplemental fig. 1B for gating strategy. 2-way ANOVA with Dunnett post-hoc. (C) Light-sheet render showing macrophage (F4/80, red) dispersed throughout an ipsilateral lumbar DRG isolated 1 day after saline or 1% carrageenan injection. See supplemental movies S1 and S2. Scale bar: 150µm. (D) Course of mechanical hyperalgesia in MM^dtr^ and control littermates (Ctr) injected with 1% carrageenan in one and saline in the other hind paw. Mixed-effects model (REML), with Sidak post-hoc. (E) Course of weight bearing of the left hind paw (as % of total body weight) in Ctr and MM^dtr^ littermates injected with 1% carrageenan in one and saline in the other hind paw. 2-way repeated-measures ANOVA, comparing WT (o) and MM^dtr^ (*) versus day 0. (F) Course of carrageenan-induced mechanical hyperalgesia in male versus female in MM^dtr^ and Ctr littermates. 2-way repeated-measures ANOVA, asterisks indicate significance between male(o)/female(*) WT vs male/female MM^dtr^ mice. (G) Course of CFA-induced mechanical hyperalgesia in Ctr and MM^dtr^ littermates. 2-way repeated-measures ANOVA, Sidak post-hoc comparing carrageenan conditions. See supplementary files for all related statistical values. Data are represented as mean±SD. Related data is available in supplemental figures 1-3.

The expression of *Il6* mRNA at day 7 was similar in the paws of MM^dtr^ and littermate control mice, indicating that carrageenan-induced inflammation was resolved in both groups of mice (Supplemental Fig. 2K). Thus, persisting mechanical hypersensitivity in monocyte/macrophage depleted MM^dtr^ mice is independent of ongoing inflammation. Coherently, the intraperitoneal injection of the immunosuppressive drug dexamethasone at day 6 after carrageenan did not relieve mechanical hypersensitivity (Fig. 2A). In contrast, dexamethasone did prevent inflammation mediated development of mechanical hypersensitivity (Fig 2B) when administered prior to the injection of carrageenan.

**Figure 2.**
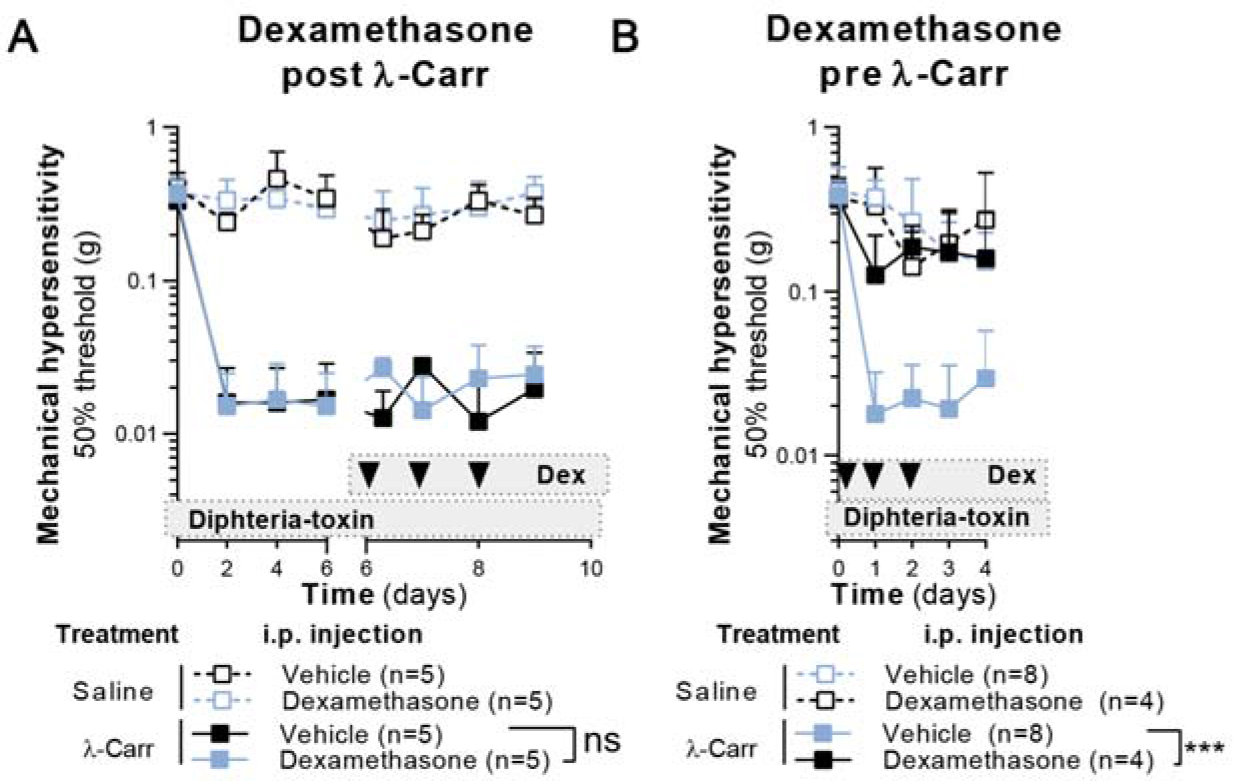
Ongoing pain in MM^dtr^ mice is not caused by ongoing inflammation. (A, B) Course of mechanical hyperalgesia in MM^dtr^ mice after i.pl. injection of 1% carrageenan or saline control in a hind paw. Mice received dexamethasone at indicated time points. 2-way repeated measure ANOVA, Sidak post-hoc.

To directly target monocytes to the DRG (Willemen et al., 2018), we intrathecally (i.t.) injected wildtype (WT) bone-marrow-derived CD115^+^ monocytes into MM^dtr^ mice (model described in Supplemental fig. 4A), which reconstituted macrophages in the ipsilateral DRG (Supplemental fig. 4B). Within hours, i.t. injection of WT monocytes sustainably rescued the defective resolution of hyperalgesia in MM^dtr^ mice (Fig. 3A and supplemental fig. 4C). The pain-resolving capacity of monocytes was independent of their origin (bone marrow or spleen; or Ly6C expression (‘classical’ Ly6C^+^ or ‘non-classical’ Ly6C^-^; supplemental figs. 4D-F). These data show that monocytes are essential to resolve inflammatory pain.

**Figure 3.**
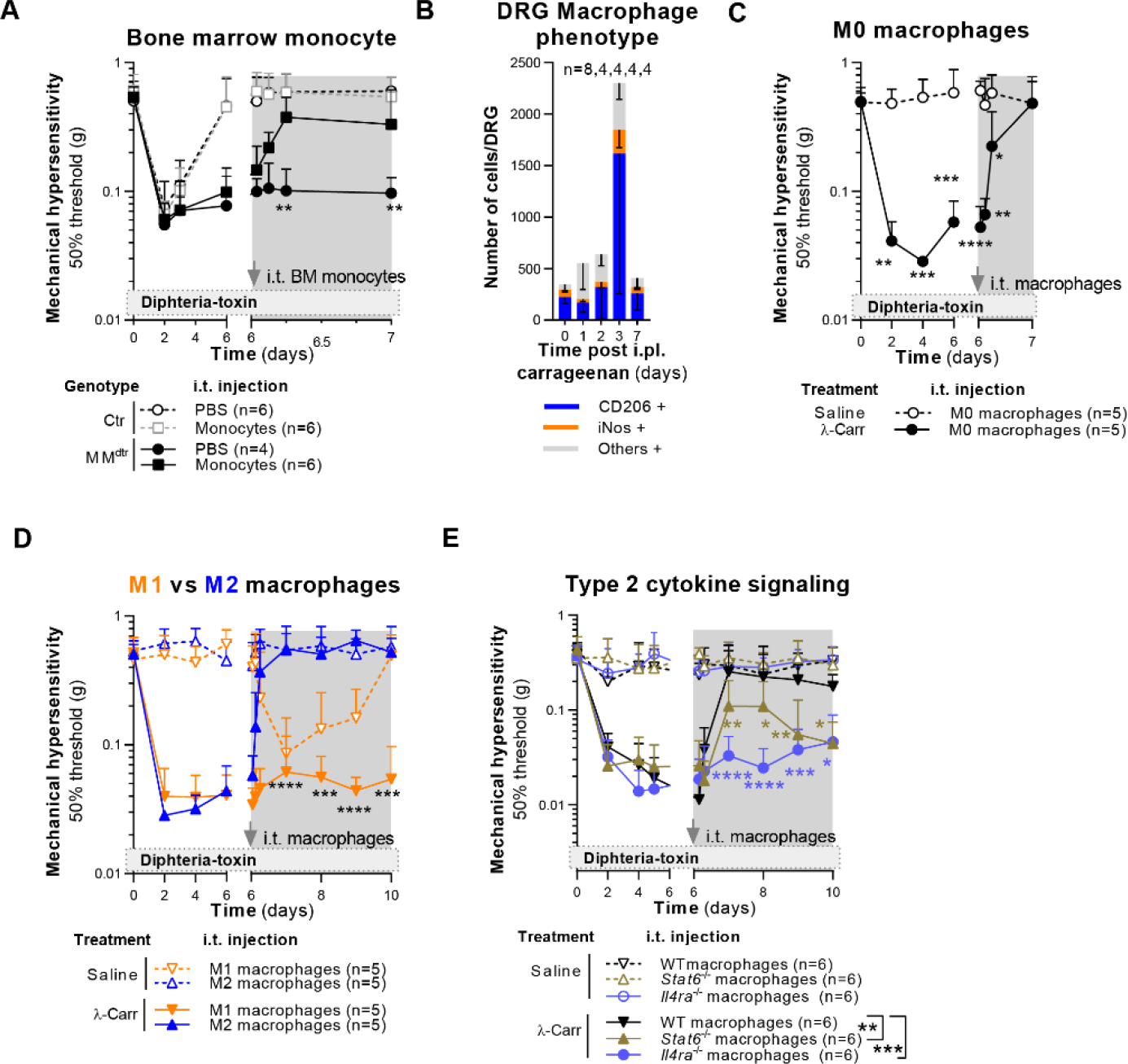
Monocytes/macrophages are required to resolve inflammatory pain. (A) Course of mechanical hyperalgesia in littermate control (Ctr) and MM^dtr^ mice after i.pl. injection of 1% carrageenan in the hind paws and i.t. injection of PBS or WT CD115^+^ monocytes. 2-way repeated measure ANOVA, Sidak post-hoc comparing MM^dtr^ conditions. (B) Phenotype of F4/80 positive macrophages in DRG of mice i.pl injected with carrageenan in the hind paw at indicated time points. Gating strategy is indicated in Fig. S1B. 2-way ANOVA with Dunnett post-hoc. (C) Course of mechanical hyperalgesia in MM^dtr^ mice after i.pl. injection of 1% carrageenan in one and saline in the other hind paw, followed by i.t. injection of M0 macrophages. 2-way repeated measures ANOVA, with Sidak post-hoc. (D) Course of mechanical hyperalgesia in MM^dtr^ mice after i.pl. injection of 1% carrageenan in one and saline in the other hind paw, followed by i.t. injection of M1 or M2 macrophages. 2-way repeated measures ANOVA, with Sidak post-hoc comparing carrageenan conditions. (E) Course of mechanical hyperalgesia in MM^dtr^ mice after i.pl. injection of 1% carrageenan in one and saline in the other hind paw, followed by i.t. injection of WT, *Stat6*^-/-^ or *Il4ra*^-/-^ macrophages. 2-way repeated measures ANOVA, with Sidak post-hoc comparing carrageenan conditions. See supplementary files for all related statistical values. Data are represented as mean±SD. Related data is available in supplemental figures 1 and 4.

Macrophages that reside in peripheral nerve tissue are different from microglia and non-nervous residing macrophages (Liang et al., 2019; Wang et al., 2020). After nerve injury, monocyte-derived macrophages engraft in the pool of resident peripheral nervous system macrophages (Ydens et al., 2020), and are programmed by vesicles secreted by sensory neuron (Simeoli et al., 2017) and can help to suppress neuropathic pain (Niehaus et al., 2021). We determined whether monocytes/macrophages that infiltrate the DRG during inflammatory pain, had an inflammatory (‘M1’) - or resolution (‘M2’)-like phenotype. At day 3, the number of CD206^+^ M2-like or tissue-repair macrophages (Bosurgi et al., 2017; Minutti et al., 2017) was increased in the DRG, whereas the number of inducible nitric oxide synthase (iNOS)^+^ M1-like or inflammatory macrophages did not significantly change (Fig. 3B). Consistent with the dominant presence of CD206^+^ M2-like macrophages, i.t. injection of *in vitro* differentiated bone-marrow derived macrophages (‘M0’, from here on referred to as ‘macrophages’) and macrophages subsequently differentiated with interleukin (IL)-4 (‘M2’) rescued resolution of hyperalgesia in MM^dtr^ mice (Figs. 3C-D, supplemental fig. 4G-H). In contrast, inflammatory macrophages differentiated with lipopolysaccharide and interferon-γ (‘M1’) induced a transient hyperalgesia in the saline treated paws and were incapable of resolving inflammatory hyperalgesia in the carrageenan treated paws (Fig. 3D and supplemental fig. 4H). In agreement with a requirement for M2-like differentiated macrophages in pain resolution, macrophages deficient for *Il4ra* or *Stat6* failed to sustainably resolve inflammatory hyperalgesia (Fig. 3E). Macrophages resolved pain through a pathway independent of neuronal IL-10 receptor (*Il10r*) signalling because *Nav1.8*^Cre^*Il10r^flox^* mice, which are deficient for the IL-10 receptor in pain-sensing sensory neurons that mediate inflammatory hyperalgesia (Abrahamsen et al., 2008), recovered normally (Supplemental fig. 4I).

Metabolically, M2 macrophages depend on oxidative phosphorylation (OxPhos), while M1 macrophages are glycolytic (Supplemental fig. 5) (Galvan-Pena and O’Neill, 2014; Van den Bossche et al., 2015). Neurons have a high metabolic demand (Misgeld and Schwarz, 2017). We previously demonstrated that a deficiency in mitochondrial function in sensory neurons prevents the resolution of inflammatory pain (Willemen et al., 2018). Moreover, in chronic pain neuronal mitochondrial functions, such as OxPhos and Ca^2+^ buffering, are impaired (Duggett et al., 2017; Hagenston and Simonetti, 2014). Indeed, oxygen consumption in DRG neurons was reduced with ∼30% during the peak of inflammatory pain and resolved preceding pain resolution at day 3 (Fig. 4A). Therefore, we posited that macrophages facilitate the restoration of a functional mitochondrial pool. Given that after ischemic stroke neurons can take up mitochondria released by adjacent astrocytes (Hayakawa et al., 2016), we hypothesized that during inflammatory pain resolution monocytes/macrophages aid neurons by supplying mitochondria.

**Figure 4.**
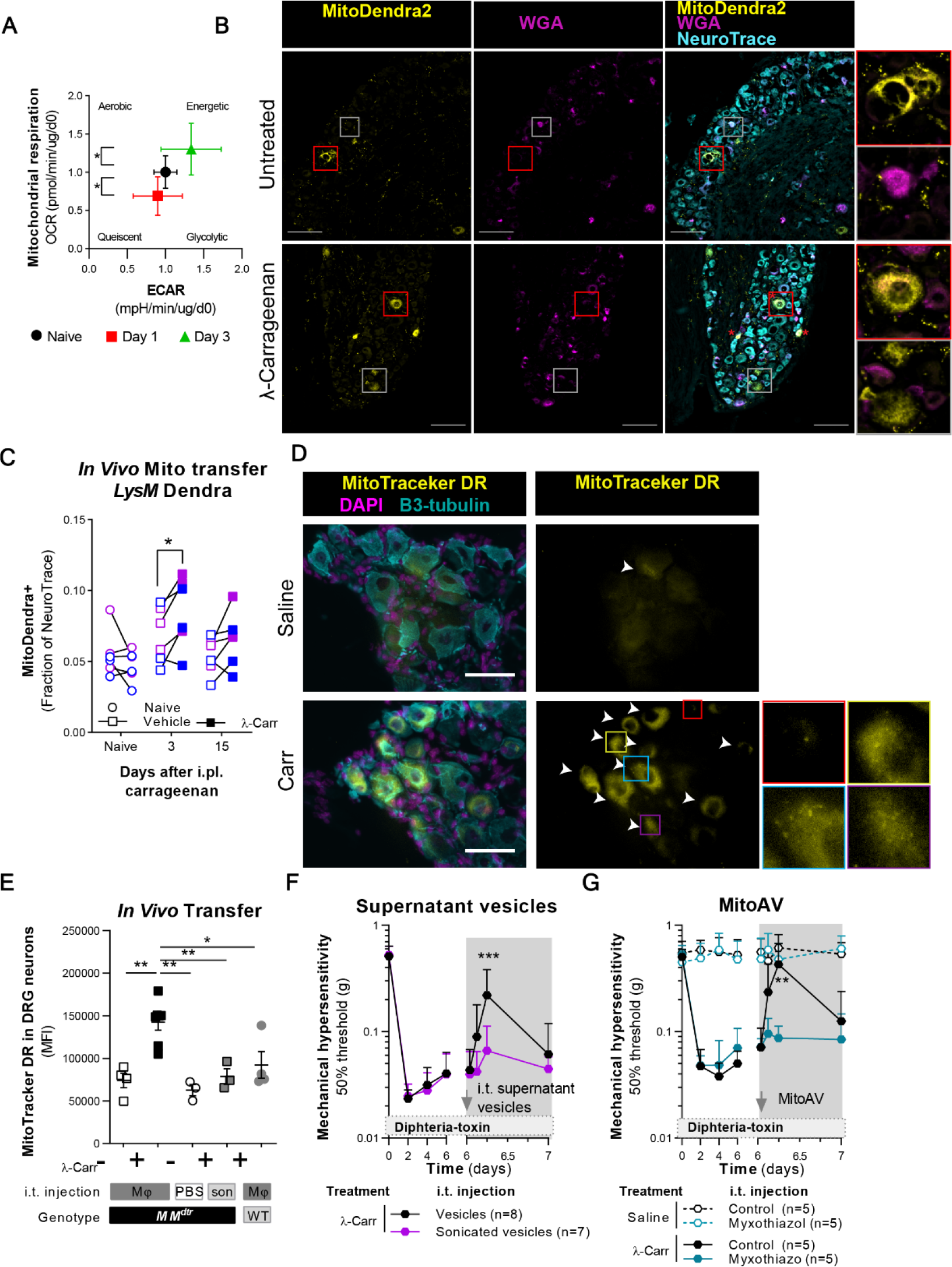
Macrophages migrate into the DRG and transfer mitochondria to neurons. (A) Basal oxygen consumption rates in sensory neuron cultures obtained from lumbar DRG isolated from mice at indicated days post carrageenan injection in the hind paw. DRG from 1 or 2 mice were pooled per experiment and divided over 3-5 wells. A total of 5 mice were assessed in 3 experiments. ANOVA with Hold-Sidak post- hoc. (B, C) (B) Example images and (C) quantification of the MitoDendra2^+^ area within the NeuroTrace^+^ area of the contra- or ipsilateral DRG of LysM^cre^-MitoDendra2^flox^ mice before, three and 15 days after carrageenan or saline injection. scale bar: 150 µm. n=6. Color coded to sex: blue; male; purple: female. Paired-t Test. (D, E) (E) Example images and (D) quantification of MTDR signal in sensory neurons in the DRG of MM^dtr^ and WT littermates. At day 6 after 1% carrageenan (ongoing pain in MM^dtr^, resolved pain in WT mice) or saline injection, we injected i.t. PBS, MTDR-labelled macrophages (Mφ), or sonicated MTDR-labelled macrophages (son). After 18h, lumbar DRG were isolated for immunofluorescence analysis and counter- stained with β3-tubulin (cyan, neurons) and DAPI (magenta, nuclei). White arrowheads indicate MTDR^+^ (yellow) neurons. Scale bar: 50µm. ANOVA with Sidak’s post-hoc. (F) Course of mechanical hyperalgesia in MM^dtr^ mice injected with carrageenan. At day 6 mice were injected i.t. with intact or sonicated macrophage-derived vesicles. 2-way ANOVA, Sidak post-hoc comparing carrageenan conditions. (G) Course of mechanical hyperalgesia in MM^dtr^ mice injected with carrageenan in the left hind paw, and saline in the right hind paw. At day 6 mice were injected i.t. with artificially generated vesicle containing mitochondria (MitoAV) with functional or myxothiazol (complex III)-inhibited mitochondria. 2-Way ANOVA with Dunnett post-hoc. See supplementary files for all related statistical values. Data are represented as mean±SD, and mean±SEM in graphs showing individual data points. Related data is available in supplemental figures 5-11.

We stained mitochondria from macrophages with MitoTracker Deep Red (MTDR) that binds covalently to mitochondrial proteins (Chazotte, 2011) and co-cultured live macrophages, or an equivalent volume of sonicated macrophages, with the neuronal cell line Neuro2a (N2A). After 2 hours, macrophage-derived MTDR^+^-mitochondria were detectable in N2A cells by flow cytometry and image stream (Supplemental figs. 6A/B). Macrophages transduced with mitochondria-targeted DsRed (mitoDsRed) also transferred mitochondria to primary sensory neurons upon co-culture *in vitro*, excluding that the signal was due to MTDR leak from macrophages to neurons (Supplemental fig. 6C). Transfer of mitochondria from macrophages to neurons also occurred *in vivo*. During the resolution of pain, the area of MitoDendra2 within the NeuroTrace area of *LysM*^Cre^-MitoDendra2^flox^ mice increased at day 3 after carrageenan injection at the ipsilateral side in males and females (Fig. 4B/C). In a separate experiment we confirmed that at day 3 the percentage of MitoDendra2 positive cells was approximately doubled at the ipsilateral side comparted to the contralateral side (Supplemental Fig. 7A/B). A proportion of the MitoDendra2^+^ neurons were also positive for the retrograde tracer WGA that was injected 2 days prior to the isolation of DRG (Fig. 4C). These data suggest that monocytes/macrophages transfer mitochondrial content to neurons innervating the paw during resolution of inflammatory pain. At day 14 we did not observe a significant difference in Mitodendra2 area between ipsi and contralateral DRG (Fig. 4C). Moreover, mitoDendra2*^flox^* mice or *LysM*^Cre^-GFP*^flox^* mice did not have any MitoDendra2 or GFP positive neurons, suggesting that MitoDentra2 positivity in neurons was not because of a leaky LysM promotor (Supplemental fig. 7C). In addition, intrathecal injection of MTDR-labelled macrophages in MM^dtr^ mice at day 6 after carrageenan injection increased the MTDR labelling (MFI and percentage) of sensory neurons from mice with persisting inflammatory hyperalgesia (Figs. 4D/E, supplemental fig. 7D), but not in control treated mice or in WT mice that had resolved inflammatory hyperalgesia. Injection of sonicated MTDR-labelled macrophages did not result in accumulation of MTDR in sensory neurons (Fig. 4E), excluding that the signal in sensory neurons was due to uptake of MTDR leaking from macrophages. We conclude that macrophages transfer mitochondria to neurons in the DGR during resolution of inflammatory pain. Using flow cytometry, we found that macrophages released CD45^+^ extracellular vesicles that stained positive for macrophage plasma membrane proteins, such as MHC class II, CD11b and CD200 Receptor 1 (CD200R), and the mitochondrial dye MTDR (Supplemental fig. 8A/B). In line with the MTDR staining in vesicles, in the supernatant of MitoDendra2^+^ macrophages ∼17% of CD45^+^ vesicles were also MitoDentra2^+^ positive (Supplemental fig. 8C). The vesicles had a broad range in size (Supplemental fig. 8D).

We hypothesized that the mitochondria-containing vesicles released by macrophages were sufficient to resolve pain. Indeed, i.t. administration of mitochondria-containing extracellular vesicles isolated from macrophage supernatant rapidly but transiently resolved inflammatory hyperalgesia in MM^dtr^ mice (Fig. 4F, supplemental fig. 8E). However, injection of extracellular vesicles that were destroyed by sonication did not affect hyperalgesia (Fig. 4F). Taken together, this suggests that intact vesicles with mitochondria, but not their individual components such as lipids and proteins, are sufficient to resolve pain. In support of the need of functional mitochondria, monocytes that have distressed mitochondria and significantly reduced mitochondrial DNA (mtDNA) content due to a heterozygous deletion of the Transcription Factor A/*Tfam* (West et al., 2015) failed to resolve inflammatory hyperalgesia in MM^dtr^ mice (Supplemental figs. 9A/B). Finally, we isolated artificial vesicles containing mitochondria from M0 macrophage cell bodies (MitoAV). MitoAV stained positive for macrophage plasma membrane markers and MTDR and had active OxPhos (Supplemental figs. 9C and 12B). Intrathecal injection of MitoAV rapidly but transiently resolved inflammatory hyperalgesia in MM^dtr^ mice (Fig. 4G and supplemental fig. 9D). In contrast, MitoAV in which oxidative phosphorylation was blocked by complex III inhibitor myxothiazol (Thierbach and Reichenbach, 1981) failed to resolve hyperalgesia (Fig. 4G and supplemental fig. 9D). Thus, vesicles secreted by or isolated from macrophages contain mitochondria and resolve inflammatory pain.

For efficient transfer of mitochondria, we hypothesized that docking of extracellular vesicles to sensory neurons is facilitated by receptor-ligand interactions. Macrophages, predominantly those with an M2 phenotype (Koning et al., 2010), and macrophages-derived extracellular vesicles expressed CD200R (Supplemental fig. 6A), while neurons are known to express its ligand CD200 (Wright et al., 2001)). In line with this reasoning, *Cd200r*^-/-^ mice failed to resolve inflammatory hyperalgesia, which persisted for at least 16 days (Fig. 5A and Supplemental fig. 10A). Place-preference conditioning with the fast-working analgesic gabapentin (Navratilova and Porreca, 2014), a drug that relieves neuropathic and inflammatory pain (Park et al., 2016; Singh et al., 1996), confirmed ongoing spontaneous pain in *Cd200r*^-/-^ mice for at least 2 weeks after carrageenan injection (Fig. 5B, supplemental fig. 10B). Of note, acute inflammation and the resolution of inflammation at the site of carrageenan injection in *Cd200r*^-/-^ mice did not differ from that of WT mice (Figs. 5C and 5D). This further supports the role of CD200R in the resolution of acute inflammatory pain and prevention of chronic pain.

**Figure 5.**
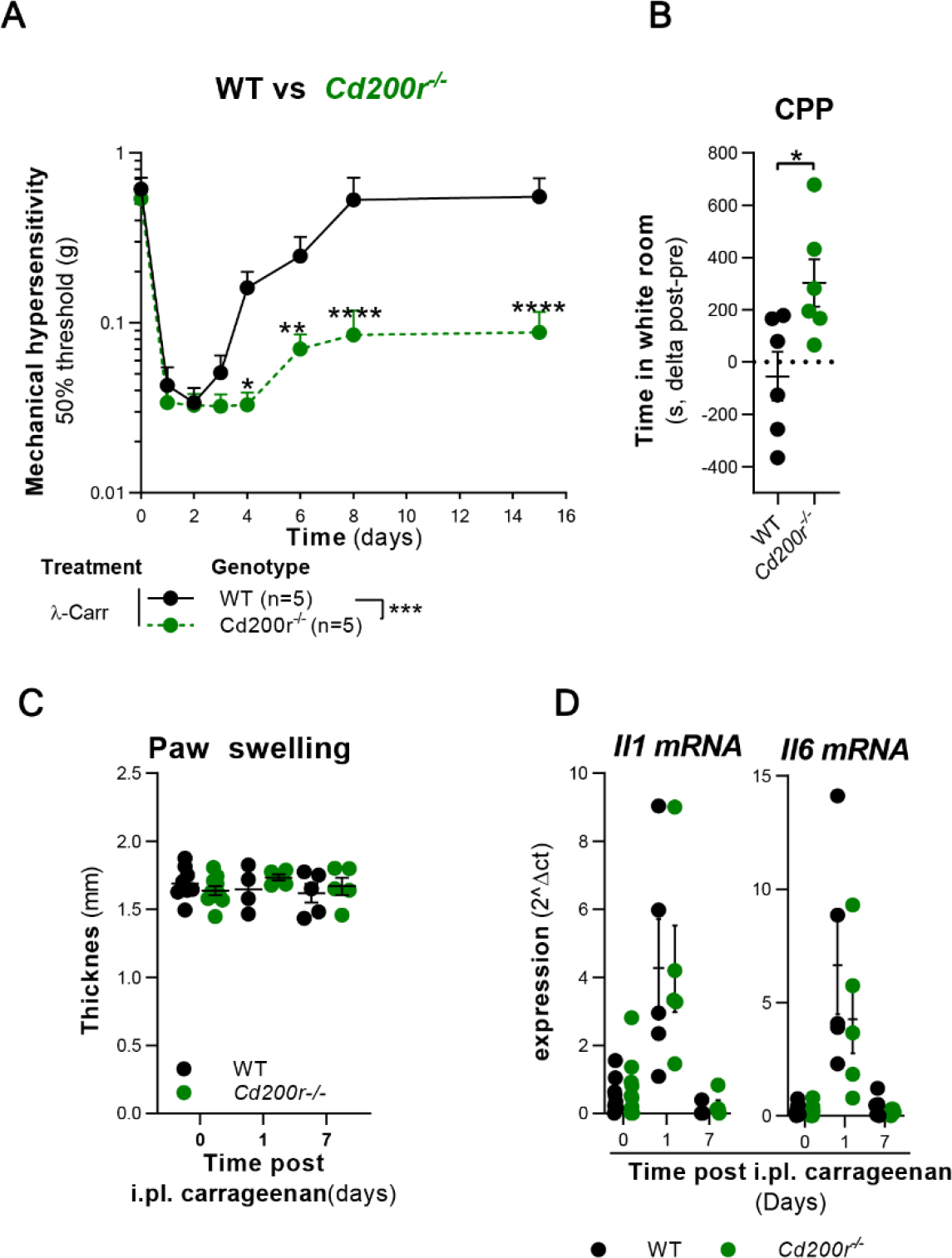
Monocytes require CD200R to resolve inflammatory pain. (A) Course of mechanical hyperalgesia in *Cd200r*^-/-^ or WT mice that were unilateral injected with 1% carrageenan in the hind paws. 2-way ANOVA, Sidak post-hoc. (B) Gabapentin-induced place preference conditioning at day 16 after unilateral 1% carrageenan injection in the hind paws. Conditioning efficiency is depicted as the difference in time (seconds, s) spent in a white room pre- and post-conditioning. Unpaired t-test. (C) Paw swelling of the carrageenan-injected paw in WT and *Cd200r*^-/-^ mice. 2-way repeated measures ANOVA, Sidak. (D) mRNA expression of cytokines *Il1* and *Il6* in the carrageenan-injected paw of WT and *Cd200r*^-/-^ mice. 2-way repeated measures ANOVA, Sidak. See supplementary files for all related statistical values. Summarized data are represented as mean±SD, and mean±SEM in graphs showing individual data points. Related data is available in supplemental figure 10.

Intrathecal injection of *Cd200r*^-/-^ monocytes did not resolve inflammatory hyperalgesia in MM^dtr^ mice (Fig 6A, supplemental fig. 11A). Consistent with these data, WT monocytes or macrophages resolved persisting inflammatory hyperalgesia in *Cd200r*^-/-^ mice, whereas injection of additional *Cd200r*^-/-^ monocytes or macrophages did not (Fig. 6B; supplemental fig. 11B). These data indicate an intrinsic defect in the pain-resolution capacity of *Cd200r*^-/-^ monocytes and macrophages independent from effects of macrophages at the site of primary inflammation.

**Figure 6.**
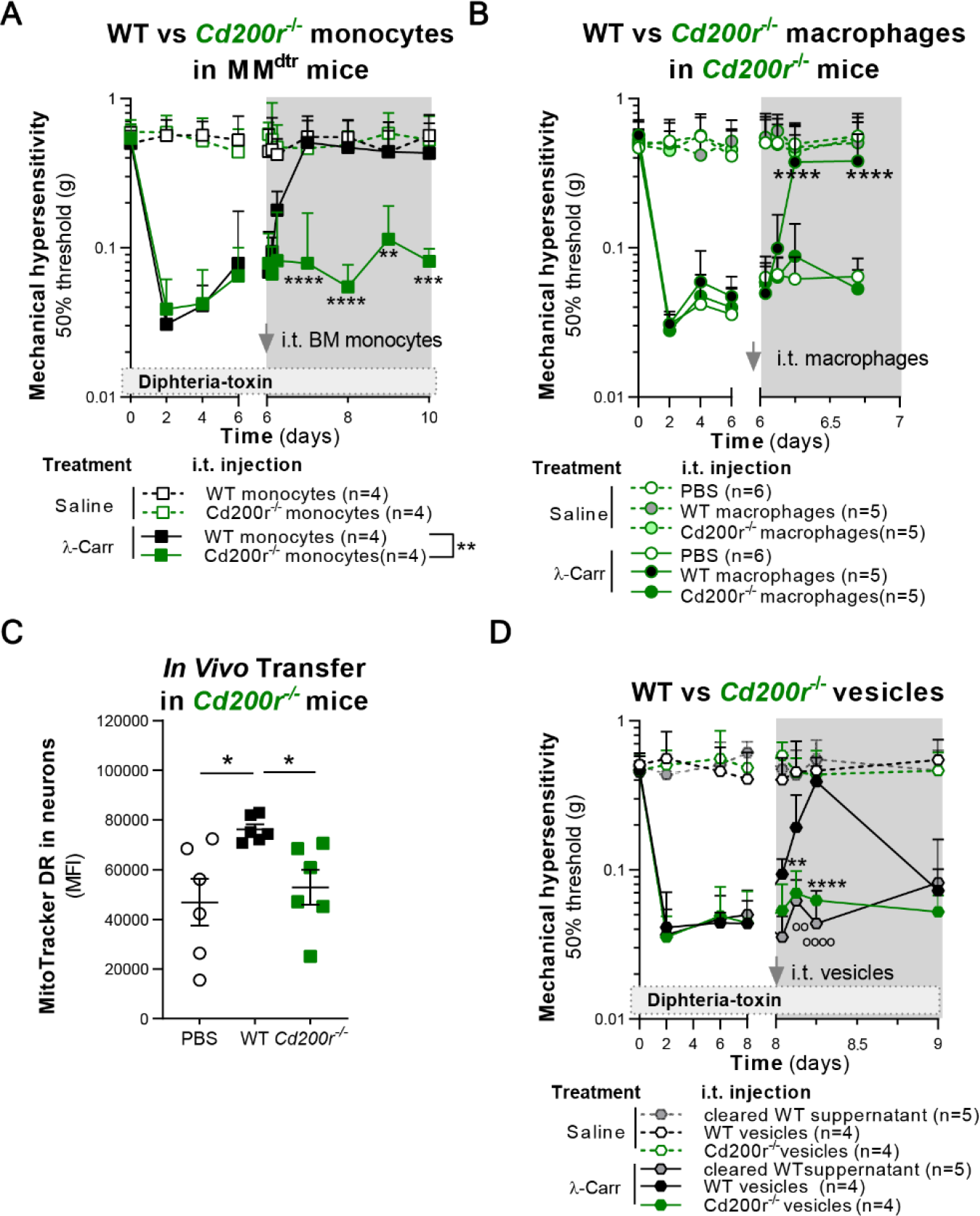
Cd200r deficient monocytes/macrophages fail to resolve inflammatory pain. (A) Course of mechanical hyperalgesia in MM^dtr^ mice that were injected unilateral with 1% carrageenan or saline. At day 6, WT or *Cd200r*^-/-^ CD115^+^ monocytes were i.t. injected. 2-way repeated measures ANOVA, Sidak post-hoc comparing carrageenan conditions. (B) Course of mechanical hyperalgesia in *Cd200r*^-/-^ mice that were injected unilateral with 1% carrageenan and saline. At day 6, WT or *Cd200r*^-/-^ macrophages were i.t. injected. 2-way repeated measures ANOVA, Sidak post-hoc comparing carrageenan conditions. (C) In vivo MTDR transfer from WT or *Cd200r*^-/-^ MTDR-labelled macrophages to DRG neurons in *Cd200r*^-/-^ mice. At day 6 after carrageenan injection, macrophages or PBS were injected i.t. and after 18h DRG were isolated and stained as described for Fig. 3D/E. ANOVA with Holm-Sidak post-hoc. (D) Course of mechanical hyperalgesia in MM^dtr^ mice injected unilateral with 1% carrageenan and saline. At day 6, vesicles isolated from WT or *Cd200r*^-/-^ macrophage culture supernatant, or the supernatant of the vesicle pellet (cleared supernatant) was i.t. injected. 2-way repeated measures ANOVA, Dunnett post-hoc comparing carrageenan conditions. See supplementary files for all related statistical values. Data are represented as mean±SD, and mean±SEM in graphs showing individual data points. Related data is available in supplemental figures 10-13.

We found no evidence for a defect in mitochondrial respiration or vesicle release in *Cd200r*^-/-^ macrophages (Supplemental figs. 12A-C) and *Cd200r*^-/-^ macrophages were normal in their capacity to migrate into the DRG and had a similar phenotype to WT macrophages (Supplemental figs. 12D-H). This suggested instead that there was a defect in mitochondrial transfer between *Cd200r*^-/-^ macrophages and neurons. MTDR-labelled mitochondria did transfer from i.t. injected MTDR-labelled WT macrophages to neurons from *Cd200r*^-/-^ mice (Fig. 6C). In contrast, *Cd200r*^-/-^ macrophages failed to transfer MTDR-labelled mitochondria to sensory neurons of *Cd200r*^-/-^ mice (Fig. 6C; supplemental fig. 13A). Thus, CD200R expression on macrophages but not on neurons is required for transfer of mitochondria from macrophages to neurons. In addition, since no MTDR staining was observed in neurons upon injection of MDTR-labelled *Cd200r^-/-^* macrophages, we exclude MTDR leaking from macrophages. Extracellular vesicles isolated from *Cd200r*^-/-^ macrophage culture supernatant or supernatant from WT vesicle-pellets did not resolve inflammatory hyperalgesia in MM^dtr^ mice (Fig. 6D; supplemental fig. 13B). Thus, CD200R expression on extracellular vesicles, and not soluble factors produced by macrophages, is required for the resolution of inflammatory pain.

CD200 is the best-known ligand for CD200R and in inflammatory models, such as arthritis, *Cd200^-/-^* and *Cd200r^-/-^* mice have a similar phenotype (Hoek et al., 2000; Simelyte et al., 2010). However, in sharp contrast to *Cd200r^-/-^* mice, *Cd200^-/-^* mice completely resolved inflammatory pain with similar kinetics to WT litter mates (Fig. 7A; supplemental fig. 14A). This suggests the involvement of an alternative CD200R ligand. In 2016, iSec1/*Gm609* was described as a CD200R ligand expressed specifically in the gut (Kojima et al., 2016). We found that iSec1*/Gm609* mRNA is also expressed in DRG, along with *CD200* mRNA (Supplemental figs. 14B/C). Using in-situ hybridization (RNAScope), we detected *iSec1/gm609* mRNA in neurons and cells surrounding neurons (Supplemental fig. 14D). Additionally, iSec1 mRNA was expressed in Advillin^+^ neurons but not in CD45^+^ cells from the DRG of Advillin^MitoDendra2^ mice (Supplemental fig. 14E).

**Figure 7.**
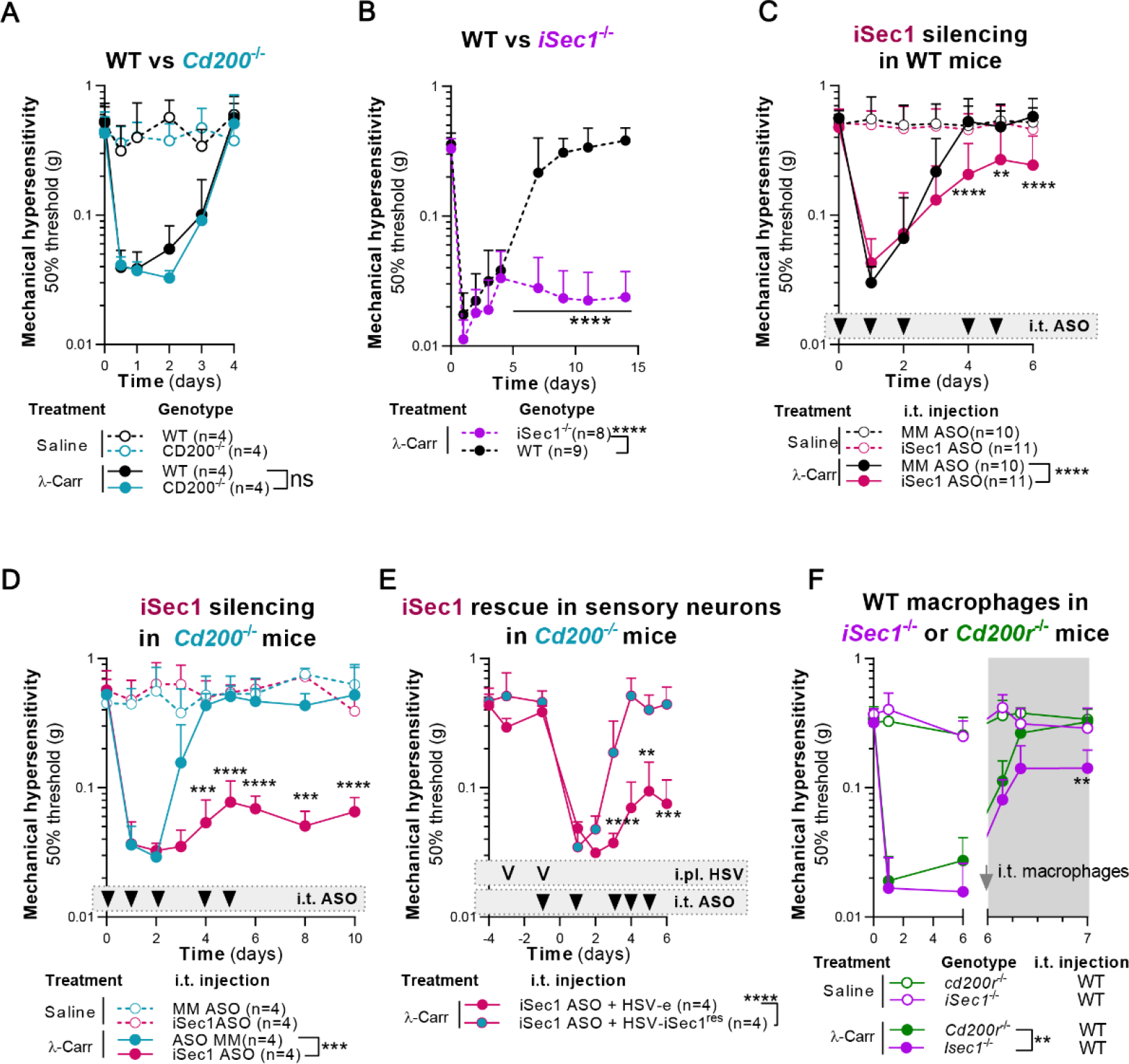
iSec1 is required for resolution of pain. (A, B) Course of carrageen-induced mechanical hyperalgesia in WT, *Cd200*^-/-^ or *iSec1*^-/-^ littermates. 2-way repeated measures ANOVA, Sidak post-hoc comparing carrageenan conditions. (C, D) Course of carrageen-induced mechanical hyperalgesia in WT (C) or *Cd200*^-/-^ (D) mice injected with mismatch (MM-ASO) or iSec1-specific antisense oligonucleotides (iSec1-ASO). 2-way repeated measures ANOVA, Sidak post-hoc comparing carrageenan conditions. (E) Course of carrageen-induced mechanical hyperalgesia in *Cd200^-/-^* mice that received intraplantar (i.pl) with HSV-e or HSV-iSec1^res^ before i.pl. carrageenan injection, treated with iSec1-specific ASO injected i.t. 2-way repeated measures ANOVA, Sidak post-hoc. (F) Course of carrageenan-induced thermal hyperalgesia in MM^dtr^ mice that received i.t. *Cd200r*^-/-^ or *iSec1*^-/-^ macrophages at day 6. 2-way repeated measures ANOVA, Sidak post-hoc. See supplementary files for all related statistical values. Data are represented as mean±SD. Related data is available in supplemental figure 14-16.

In contrast to *Cd200*^-/-^ mice, *iSec1*^-/-^ mice failed to resolve pain (Fig. 7B, supplemental figs. 15A/B). Coherently, silencing iSec1 mRNA expression in the DRG of WT mice by repetitive intrathecal injections of *iSec1/Gm609* targeting antisense oligodeoxynucleotides (ASO) (Fig. 7C and supplemental fig. 16C/D) partially prevented resolution of inflammatory hyperalgesia (Fig. 7D; Supplemental fig. 15E) (Lai et al., 1996). In *Cd200*^-/-^ mice, i.t. injections of *iSec1/Gm609*- ASO completely prevented the resolution of hyperalgesia (Fig. 7D; supplemental fig.15F). Next, we used intraplantar injection of Herpes Simplex Virus (HSV) to specifically target sensory neurons innervating the inflamed area for exogenous expression of iSec1 (Willemen et al., 2018). Expression of *iSec1/gm609* that was mutated to resist ASO treatment in sensory neurons (HSV-iSec1^res^, Supplemental fig. 15G) completely rescued the ability of iSec1*/Gm609*-ASO treated *Cd200*^-/-^ mice to resolve pain, while an empty vector (HSV-e) did not (Fig. 7G; Supplemental fig. 15H). We further evaluated whether the inability of *Isec1*^-/-^ mice to resolve pain is rescued by injection of WT macrophages. In coherence with expression patterns (Supplemental fig. 14), WT macrophages do not fully resolve pain in *iSec1*^-/-^ mice (Fig. 7F). We conclude that monocyte/macrophage expression of CD200R and sensory neuron expression of the ligand iSec1 is required *in vivo* to resolve inflammatory pain.

## Discussion

We identified a previously unappreciated role for macrophages which transfer mitochondria to somata of sensory neurons to resolve inflammatory pain. Previous studies showed that respiratory competent mitochondria are present in human whole blood (Al Amir Dache et al., 2020), and that tissue-resident cells can transfer mitochondria (Hayakawa et al., 2016; Moschoi et al., 2016). We now show that non-tissue resident monocytes are recruited to the DRG, acquire a M2/tissue-repair like phenotype and transfer mitochondria to sensory neurons via a CD200R:iSec1 interaction to facilitate resolution of inflammatory pain. In contrast to M2 macrophages, inflammatory M1 macrophages induced pain. Thus, a DRG-milieu that skews local macrophages towards a M1 phenotype could contribute to the development of chronic pain (Barclay et al., 2007; Liu et al., 2000; Mert et al., 2009; Zhang et al., 2016).

Previous studies have implicated macrophages in resolution of inflammatory pain at the site of the primary inflammatory insult by secretion of IL-10 (da Silva et al., 2015; Klionsky et al., 2016), or by clearance of the inflammatory agent zymosan (Bang et al., 2018). We now show that macrophages are necessary to resolve pain distant from the primary inflammatory insult and independent of ongoing (systemic) inflammation. Importantly, resolution of inflammatory pain was independent of IL-10 receptor signalling in sensory neurons, excluding a direct effect of IL- 10 on neurons in resolution of inflammatory pain.

Our data show that transfer of mitochondria by macrophages in the DRG is required for resolution of inflammatory pain. However, it is possible that macrophages have additional roles in other areas of the nervous system, including distant nerves or nerve endings. Indeed, we also observed an increase of macrophages in the spinal cord during inflammatory pain. However, after i.t. injection of mononuclear cells in mice with L5-spinal nerve injury, mononuclear cells integrate into the lumbar DRG but not into the spinal cord (Takamura et al., 2020). Moreover, intrathecal injection of monocytes/macrophages required 233 times fewer cells compared to intravenous injection (Willemen et al., 2014). This suggests that resolution of pain is regulated in neuronal tissue in, or in close proximity of, the DRG. We cannot exclude other cells to contribute to resolution of inflammatory pain. For example, *Csf1r/LysM* negative macrophages at the site of the primary inflammatory insult or satellite glial cells in the DRG that surround the soma of sensory neurons may play additional roles (Bang et al., 2018; da Silva et al., 2015; Klionsky et al., 2016; Niehaus et al., 2021).

The classical nomenclature of macrophages differentiation (M1 vs M2) does not fully reflect the enormous heterogeneity macrophages can adopt in various tissues (Niehaus et al., 2021; Wang et al., 2020; Ydens et al., 2020). We here show that expression of the M2-like marker CD206 is increased in macrophages during resolution of pain. We confirmed that macrophages require differentiation towards the ‘M2-spectrum’ to resolve pain using *Stat6*^-/-^ and *Il4ra*^-/-^ macrophages. In support, CD206^+^ macrophages control neuropathic pain (Niehaus et al., 2021). In depth analysis of the exact type of M2 macrophage will support further delineation of the exact requirements of macrophages to resolve inflammatory pain.

Previously intercellular transfer of cellular organelles was shown (Torralba et al., 2016). We show with the use of chemical and genetic approaches that macrophages transfer mitochondria to neurons *in vitro* and *in vivo*. These experiments indicated that ∼3-20% of neurons acquired mitochondria from macrophages during inflammation, which is concordant with the percentage of somata of neurons in the DRG that innervate the hind paw (da Silva Serra et al., 2016; Emery et al., 2016; Lu et al., 2001). Intriguingly, also under non-inflammatory conditions, we observed that neurons acquire some mitochondria, yet to a lesser extent than after induction of inflammatory pain. It remains to be elucidated what determines which neurons acquire macrophage-derived mitochondria.

Various chronic pain states, such as chemotherapy-induced pain and neuropathic pain caused by trauma or diabetes, are associated with mitochondrial defects (Fidanboylu et al., 2011; Flatters, 2015; Joseph and Levine, 2006; Lim et al., 2015), and 67% of patients with mitochondrial disease have chronic pain (van den Ameele et al., 2020). We show here that oxidative phosphorylation is reduced during the peak of transient inflammatory pain but is restored when inflammatory hyperalgesia resolves in WT mice. However, we found no difference in DRG OCR between WT, *Cd200r*^-/-^ and *iSec1*^-/-^ mice at day 14 after carrageenan injection (Supplemental fig. 16). Hence, restoration of OCR in DRG is not sufficient to enable resolution programs in sensory neurons. We postulate that other mitochondrial functions, like Ca^2+^ buffering, are needed to resolve pain and require help from DRG macrophages.

Given that the injection of isolated extracellular vesicles transiently resolves pain, a more durable resolution of pain requires a prolonged flux of mitochondria and/or additional signals from intact macrophages. These mitochondria could replace mitochondria in neurons that have incurred mitochondrial damage. Future work should assess how exactly neuronal mitochondrial homeostasis is restored by macrophage-derived mitochondria.

Although diverse structures, such as tunnelling nanotubes, can mediate intercellular mitochondrial transfer (Jackson et al., 2016; Terashima et al., 2005), our data show that mitochondria containing vesicles are sufficient to resolve pain. We cannot exclude that other mechanisms such as tunnelling nanotubes or cytoplasmic fusions may contribute to the observed transfer of mitochondria to sensory neurons.

CD200 has long been thought of as the only ligand for CD200R. Although previous studies implicate CD200 as a checkpoint for microglia cell activation in neuropathic pain by ligating microglial CD200R (Hernangomez et al., 2016; Zhang et al., 2011), we show that *Cd200^-/-^* mice fully resolve inflammatory pain. In contrast, iSec1^-/-^ mice do not resolve inflammatory pain. *iSec1*/*gm607* mRNA expression was low but detectable in sensory neurons but we could not confirm iSec1 protein expression due to lack of available antibodies. However, by HSV-mediated reconstitution specifically in sensory neurons, we did demonstrate that sensory neuron- iSec1 expression is required to resolve inflammatory pain. Of note, iSec1/*gm609* knockdown did have a greater effect on pain resolution in *Cd200^-/-^* mice than in WT mice, suggesting that the function of these ligands could be partially redundant. CD200R signalling induces STAT6- mediated expression of FOXP3 in microglia (Yi et al., 2016). *Stat6*^-/-^ macrophages in MM^dtr^ mice, but also WT macrophages in *iSec1*^-/-^ mice partially resolved inflammatory pain. These data could suggest that iSec1:CD200R signalling to STAT6 in macrophages may also contribute to program macrophages to resolve inflammatory pain.

Why would sensory neurons require external help to restore the integrity of their mitochondrial network? Sensory neurons face unique challenges in maintaining a functional mitochondrial network because of their exceptional architecture and their intense demand for energy to support energetically expensive processes such as resting potentials, firing action potentials and calcium signalling (Misgeld and Schwarz, 2017; Vergara et al., 2019). Stressed neurons, e.g. during inflammatory pain, turn to anabolic metabolism (Jha et al., 2015). In the face of this high energy demand during stress, an energy consuming process such as rebuilding the mitochondrial network would not be favourable. Moreover, maintaining an excess mitochondrial pool that is capable of handing the stress of inflammatory pain would come at a fitness cost. Thus, we propose that it would be more energy favourable for the organism to fulfil peak energy demands in indispensable sensory neurons by mitochondrial transfer from dispensable monocytes/macrophages.

Together, our data show that pain is actively resolved by an interaction between the immune and neuronal systems that is separate from the cessation of inflammation within the peripheral tissue. Novel therapeutic strategies to resolve chronic pain may focus on the restoration of mitochondrial homeostasis in neurons or on enhancing the transfer of mitochondria from macrophages.

## Supporting information

related statistical values

## Acknowledgments

The first three authors (RR, MV, HW) contributed equally to this manuscript and are allowed to re-order the sequence of the first authors to represent this if needed in, for example, their CV.

We would like to thank R.J. Soberman (Massachusetts General Hospital and Harvard Medical School) for sharing *Cd200r*^-/-^ mice; W. Muller (University of Manchester) for providing us with the *LysM*^cre^ x *Il10r*^flox^ mice; B. Chan and J. Allen (University of Manchester) for Il4ra^-/-^ bone marrow, Gerald Shadel and Zheng Wu (Salk Institute) for providing *Tfam*^+/-^ bone marrow; F. Baixauli and E. Pearce (both Max Planck Institute of Immunobiology and Epigenetics) for MitoDendra2 expressing bone marrow; Y. Adolfs (UMC Utrecht) for assisting with lightsheet microscopy; P. Vader (UMC Utrecht) for help with particle analysis, L. van Vliet, M. Bruel, and J. Raemakers (all UMC Utrecht) for their help with the analysis of immunofluorescence pictures; B. Burgering (UMC Utrecht) and S. Kaech (Yale University) for access to the seahorse machine; R. Medzhitov (Yale University) and lab members for discussions; and M. Pascha (UMC Utrecht) for help with the macrophages cultures and setting up flow-cytometry protocols.

## Author contributions

Conceptualization (LM, MV, NE), Methodology (NE, LM, MV, HW, RR, JP, RL, TK, HK), Formal Analysis (MV, NE, HW, RR, JP), Investigation (LM, NE, MN, RR, HW, MVos, WKH, MV, SV, RL), Resources (RJP), Writing original draft (MV, NE, RR), Writing reviewing and editing (MV, NE, LM, RR, JP, HW, WKH), Visualization (MV, NE, RR), Supervision (MV, HW, NE, LM, RR), Project Administration (NE, LM), Funding Acquisition (HW, MV, NE, LM).

## Supplementary Materials

Materials and Methods

Figures S1-S16

Movies S1-S2

## Method section

### Lead contact

Further information and requests for resources and reagents should be directed to and will be fulfilled by the Lead Contact, Niels Eijkelkamp.

### Data availability

All data is available in the manuscript or supplementary materials. Raw data and materials will be made available upon request. Some materials used in this manuscript are subject to Material Transfer Agreement (MTA).

### Materials and Methods

#### Animals

All experiments were performed in accordance with international guidelines and approved by the experimental animal committee of University Medical Center Utrecht (license number: 2014.I.08.059) or by the local experimental animal welfare body and the national Central Authority for Scientific Procedures on Animals (CCD, license number AVD115002015323). Adult (age 8–15 weeks) male and female C57Bl/6J, *Lysm*^Cre/+^ x *Csf1r* ^LsL-DTR/+^ (Jackson laboratories #024046)(MM^dtr^), *Nav1.8*^Cre^*Il10r*^flox^, *Cd200*^-/-^ (Hoek et al., 2000), *Cd200r*^-/-^ (Soberman et al., 2012), *Lysm*^Cre^ x PhAM^flox^ (LysM^MitoDendra2^, Jackson laboratories #18397) mice in a CD57Bl/6 background were used and maintained in the animal facility of the University of Utrecht. *Nav1.8*^cre^ mice were kindly donated by Dr. Wood (University College London, UK). *Il10r*^flox^ mice were backcrossed from *Lysm*^cre^*Il10r*^flox^ mice that were kindly donated to us by Dr. Muller (University of Manchester, UK). *Cd200r*^-/-^ were kindly donated by Dr. R.J. Soberman (Harvard Medical School).

*Isec1* deficient mice: A B6N mouse BAC clone containing mouse *iSEC1* and *iSEC2* genes (RBD 7573) was provided by the RIKEN BRC through the National Bio-Resource Project of the MEXT Japan. The targeting vector for replacing the exon 2, intron 2 and exon 3 of the iSEC1 gene with the loxP-PGK-NeomycinR-loxP cassette was constructed by Red/ET recombination method according to the manufacturer’s instruction (Gene Bridges, Heidelberg, Germany). ES cell engineering and generation of knock-out mice were performed by UNITECH Co., Ltd. (Chiba, Japan) as a custom order. The floxed NeomycinR-cassette was excised by crossing with CAG-Cre mice.

Mice were housed in groups under a 12h:12h light:dark cycle, with food and water available *ad libitum*. The cages contained environmental enrichments including tissue papers and shelter. To minimize bias, animals were randomly assigned to the different groups prior to the start of experiment, and experimenters were blinded for the treatments and genotypes. For all genetic backgrounds the littermates of heterozygous breeding’s with the specified genotype were used. In all experiments, we used both sexes based on availability to correctly control for genetic background and age.

#### Transient inflammatory pain models

For the carrageenan model, mice received an intraplantar injection of 5 μ λ arrageenan (1% w/v, Sigma-Aldrich) in one or both hind paws (Wang et al., 2013). For the transient complete Freund’s adjuvant (CFA) model, mice received intraplantar injection of 2.5 μ mix of 1:1 saline and CFA (Sigma-Aldrich). In experiments where mice received a unilateral intraplantar injection, latency times and 50% thresholds of each paw was considered as an independent measure, while in experiments with bilateral intraplantar injection the average of the left and right paw were considered as an independent measure.

We performed both uni- and bilateral models to facilitated internal controls within one animal with unilateral models, or obtain more tissue per mouse in bilateral models. For both models, we have permission in our licenses (see above, under Animals).

#### Pain behavioral tests

Prior to the start of experiments, mice were first acclimatized to the testing environment by placing the mice in the in test environment for at least 3 times for 1 hour, 1-2 weeks before starting of the experiments. Behavioral assays are performed at the same time point at the day per individual experiment, preferably between 9-15h. At least 3 baselines measures are performed at different days one week before starting the experiment, with the last one at the day of the start of the experiment. Experiments were performed in the same room and same test setup over the duration of the experiment. Experimenters were well trained to perform the assays and were blinded for the experimental groups and genotypes. Experimenters did not wear any type of perfume or musk-based deodorants at the time of assessing behavior. Mice were put into the testing environment (von Frey and Hargreaves) at least 30 minutes before performing measurements with the experimenter present in the same room. The outside of the plexiglass chambers (Size: 13.0 × 25.0 × 15.0 cm) used for the von Frey and Hargreaves setup were covered to avoid that mice could see the experimenters.

Mechanical thresholds were assessed in both hind paws using the von Frey test (Stoelting, Wood Dale, IL, US) with the up-and-down method to determine the 50% threshold(Chaplan et al., 1994; Eijkelkamp et al., 2010). In short, von Frey filaments were applied for 5 seconds to the plantar surface of the paw. After applying the first filament (0.4 g), in case of a non-response the next filament with a higher force was used. In case of a response, the next lower force filament is used. A minimum of 30 seconds between the application of filaments was taken. Four readings were obtained after the first change of direction. Experimenters applied the von Frey filaments perpendicularly, smoothly, without moving the filaments horizontally during application. If the respond was ambiguous, the experimenter waited for a minute and reproved the mouse. Heat withdrawal latency times were determined in both hind paws using the Hargreaves test (IITC Life Science)(Eijkelkamp et al., 2010; Hargreaves et al., 1988). Briefly, the Hargreaves test was carried out using a perspex enclosure on a heated (32°C) glass bottom enclosure. A radiant heat source is positioned underneath the animals and aimed at the plantar surface of the hind paw. The time taken to withdraw from the heat stimulus is recorded as the withdrawal latency. Each paw is measured at least 3 times with at least 30 seconds between measurements. The intensity of the light source was adjusted to produce withdrawal latencies of 8-10 seconds in naive C57Bl/6 mice, with a pre-determined cut off at 20 seconds to prevent tissue damage.

Changes in weight bearing were evaluated using the dynamic weight bearing (DWB) apparatus (Bioseb, Vitrolles, France)(Prado et al., 2018) and the following parameters were used for the analysis. (i) Low weight threshold of 0.5g, (ii) High weight threshold of 1g, (iii) Surface threshold of 2 cells, (iv) Minimum 5 images (0.5 seconds) for stable segment detections. The device consists of a small Plexiglas chamber (11.0L×L19.7L×L11.0Lcm) with a sensor mat containing pressure transducers. The system records the average weight that each limb exerts on the floor. Mice were placed in the chamber at least 1 minute prior to starting the measurements and were allowed to move freely within the chamber. Each mouse was recorder for period of 5L inutes. A camera at the top of the enclosure was used to record all movements to ensure accurate validation of the position of the mouse by the experimenter. The weight bearing of the affected paw (ipsilateral paw) was expressed as percentage of body weight.

Spontaneous pain was assessed by the conditioned place preference test (CPP, Stoelting, Wood Dale) as described previously (Navratilova and Porreca, 2014; Park et al., 2013). The CPP apparatus consisted of 2 visually distinct chambers (18.0 × 20.0 cm) connect by a smaller neutral chamber (10.0 × 20.0 cm). The two chambers had differing visual cues, one being darker (‘dark chamber’) and one brighter (‘bright chamber’). Prior to the start of CPP, mice were acclimatized to the setup. At the baseline measurement day, the animals were placed in the center hallway with free access to both chambers. The time spent in each chamber was recorded for 15 min to determine preconditioning baseline. In case a mouse preferred the bright chamber they were excluded from further testing. Mice underwent three trial conditionings the following 3 day. In the morning of the conditioning day, the animals received intraperitoneal saline for 20 min in the preferred ‘dark’ chamber, with no access to the other chambers. In the afternoon approximately 4 hours later, mice received intraperitoneal gabapentin (100 mg/kg, Sigma-Aldrich) and were placed in the non-preferred ‘bright’ chamber for 20 min, with no access to the other chambers. After the three trial conditioning animals were placed back into the hallway between the CPP chambers with free access to all chambers for 15 min. The time spent in each chamber was again recorded. CPP was calculated by subtracting the mean time spent in the bright chamber during preconditioning (day 1) from the time spent in the bright chamber after 3 days of conditioning (day 5).

#### Depletion of monocytes and macrophages

To deplete monocytes and macrophages in vivo, MM^dtr^ mice received a first intraperitoneal injection of 20 ng/g body weight diphtheria toxin (DT) (Sigma-Aldrich) followed by daily intraperitoneal injections of 4 ng/g body weight on all subsequent days as described previously (Schreiber et al., 2013).

#### Monocyte isolation and in vitro differentiation

To obtain bone marrow-derived monocytes, the tibia and femur bones were flushed with phosphate buffered saline (PBS). For spleen-derived monocytes, spleens were mechanically excised and minced in PBS and passed through a 70 μm cell strainer (Corning). After erythrocyte lysis with RBC lysis buffer (eBioscience), cells were centrifuged on a Ficoll density gradient (GE Healthcare) for 22 min at 1100RCF at 22°C to obtain mononuclear cells. Finally, CD115^+^ monocytes were isolated with biotin labeled anti-CD115 antibody and streptavidin-coupled magnetic beads according to the manufacturer’s instructions (Miltenyi Biotec).

To obtain classical (Ly6c^hi^) or non-classical (Ly6c^low^) monocytes for adoptive transfer, monocytes were FACS sorted (FACS Aria III, BD) using CD115 and Ly6c antibodies (see subheading ‘antibodies’). *Tfam^+/-^* splenocytes and bone-marrow cells were isolated at Yale School of Medicine and shipped frozen in 10% DMSO to the Netherlands.

For monocyte-derived macrophage generation, 10×10^6^ bone-marrow cells were seeded in a 75 cm^2^ non-treated tissue culture flasks (VWR, Radnor, PA) for 7 days in macrophage medium (High-glucose Dulbecco’s Modified Eagle medium (DMEM; Cat# 31966-021, Gibco) and DMEM/F12 (Cat#31331-028, Gibco) (1:1), supplemented with 30% L929 cell-conditioned medium (see cell lines and primary cell cultures), 10% fetal bovine serum (FBS; Cat# 10270- 106, Gibco), 1% Penicillin/Streptomycin (Gibco) and 1% L-Glutamine (200 mM, ThermoFisher).

To polarize macrophages toward M1- or M2-like macrophages, cells were stimulated with 20 ng/ml IFNγ and 100 ng/ml LPS, or 20 ng/ml of IL-4 for 24 hours, respectively.

#### Adoptive Transfer of monocytes and macrophages

Cells were injected intrathecal (30.000 cells/5 µl per mouse) under light isoflurane anesthesia as described previously(Eijkelkamp et al., 2010; Hylden and Wilcox, 1980). For some experiments, 2 ×10^6^ macrophages were labeled with 20nM of MitoTracker DeepRed FM (MTDR; Thermo Fisher Scientific) in 400µl macrophage-medium for 30 minutes at 37°C, followed by 3 washes before intrathecal injection of 30.000 cells. For some experiments, cells were sonicated with a sonicator (soniprep150, MSE, UK) at a frequency of 23 kHz, 3 times for 15 seconds on ice before injection.

#### Isolation and injection of extracellular vesicles

To isolate macrophages-derived extracellular vesicles, 10×10^6^ macrophages were cultured in a 75 cm^2^ flask the day prior to the isolation of vesicles. The supernatant from 10 flasks were collected and centrifuged at 2000RCF for 10 minutes to pellet large debris(Hayakawa et al., 2016). The supernatant was divided in 1.5 ml Eppendorf tubes followed by a centrifugation at 17,000RCF for 30 minutes at 4°C. Supernatant was discarded, except the last 50 µl to resuspend the pellets and pool them together. The pooled supernatant was centrifuged at 17.000RCF for 30 minutes. The final supernatant was collected as ‘cleared supernatant’, and the pellet was resuspended in 100 µl PBS and injected intrathecally (5 µl per mouse).

To destroy vesicles, supernatant vesicles were sonicated 3 times at a frequency of 23 kHz for 15 seconds on ice before injection.

#### NTA analysis of extra cellular vesicles

For analysis of extracellular vesicles, 4 million macrophages were seeded in a T75 flask and cultured for 4 days, then they were washed and 7 ml of plain Opti-Mem (Gibco, 31985062) was added for an additional 24h. As control, mock-treated medium that was treated identically except for no macrophages had been present. Supernatant was harvested, spun down at 2000RCF to remove cell debris, followed by centrifugation at 17.000RCF to pellet the vesicles. The pellets were resuspended in PBS and measured in a Nanosight NS500 analyser (Melvern Intruments) equipped with a 405 nm laser. NTA analysis was acquired in NTA 3.3 Dev Build 3.3.104, the camera level was set at 10, and detection threshold was set at 5, all other settings were automated.

#### MitoAV: isolation and injection of mitochondria

As described before (Iuso et al., 2017), in brief: 10×10^6^ macrophages were seeded in a 75 cm^2^ flask the day prior to the isolation of mitochondria. To inhibit mitochondrial oxidative phosphorylation, macrophages were cultured for 10 minutes with 1μM myxothiazol (Sigma-Aldrich). Subsequently, macrophages were detached, and pelleted macrophages were solved in MIB buffer (210 mM D-mannitol, 70 mM Sucrose, 5 mM HEPES, 1 mM EGTA, and 0.5% (w/v) fatty acid-free BSA, pH 7.2), transferred into a glass tube, and disrupted by 30 strokes with a homogenizer. After centrifugation (600RCF, 10 min, 4°C), the supernatant is collected into a new tube and centrifuged at 8000RCF (10 min 4°C). Pellet is washed once with MIB and once with PBS. Finally, the pellet, containing the mitochondria, is resuspended in 200 µl of PBS and injected intrathecally (5 µl per mouse).

#### Cell lines and primary cell cultures

Mouse neuroblastoma N2A cells (ATCC) were kept in cell culture-medium: DMEM (Cat# 31966-021, Gibco) plus 10% FBS (Cat# 10270-106, Gibco) and 1% Penicillin/Streptomycin (Gibco) and 1% L-Glutamine (200 mM, ThermoFisher).

DRG were collected, and subsequently digested in an enzyme mixture containing Ca2^+^- and Mg2^+^-free HBSS, 5□M HEPES, 10□M glucose, collagenase type XI (5 mg/ml) and dispase (10 mg/ml) for 40 min before mechanical trituration in DMEM+10% heat-inactivated fetal bovine serum. Cells were centrifuged for 5 min at 140 RCF, resuspended in DMEM containing 4.5 g/l glucose, 4□M L-glutamine, 110□g□/l sodium pyruvate, 10% fetal bovine serum, 1% penicillin–streptomycin (10,000 i.u./ml), 1% glutamax, 125 ng/ml nerve growth factor, and plated on poly-l-lysine- (0.01□g/ml) and laminin- (0.02□g/ml) coated 35-mm dishes. Neurons were used 24 h after plating.

To obtain L929 cell-conditioned medium, 10×10^6^ L929 cells were seeded in a 75 cm^2^ flask with cell culture-medium supplemented with 1% non-essential amino acids (Sigma-Aldrich) for a week. L929 cells were passaged to a 162 cm^2^ flask with 50 ml medium and after a week the supernatants were collected and filtered through a 0.2- μm filter and stored at −20°C (L929-drived M-CSF).

#### Over-expression and cloning

We amplified CD200 cDNA derived from DRG (mCD200-BamHI-fwd: TAAGCAGGATCCGCCGCCACCATGGGCAGTCTGGTATTCAG; mCD200-SalI-rev: TGCTTAGTCGACTCATTATTTCATTCTTTGCATCC; mCD200-NotI-rev:TGCTTAGCGGCCGCTCATTATTTCATTCTTTGCATCC) and ligated the PCR product into pMXc after digestion with BamHI and ApoI, or into pLenti-MP2 (Enomoto et al., 2013) after digestion with BamHI and SalI. All cDNA inserts were verified using sanger sequencing.

Standard transduction protocols were performed to generate stable CD200 and iSec1 expression in N2A using pMX-iSec1-IRES-GFP(Kojima et al., 2016) and pLenti-MP2-CD200. MitoDsRed expression in macrophages were made by standard transduction protocols using pLV-MitoDsRed(Kitay et al., 2013).

We generated a bicistronic herpes simplex virus (HSV) construct by cloning iSec-Flag, under control of the α promotor and GFP under control of the α promoter (HSV-iSec). We used sewing-PCR to introduce silent mutations in iSec1-Flag to resist antisense oligonucleotide-mediated knockdown (see table below for sequence of primers). The first PCR products were made by combining primers ‘*iSec1-FLAG forward’* with ‘iSec1^resASO1^_mid_reverse’, and ‘*iSec1^resASO1^_mid_forward*’ with ‘*iSec1^resASO2^-reverse*’. The right length products were excised from agarose gel, purified and combined in a next PCR using primers ‘*iSec1-FLAG forward’* with ‘*iSec1^resASO2^-reverse*’. The resulting iSec1^res^ was digested with HINDIII and purified from agarose gel and ligated into HSV as described before (Roy et al., 2002), and validated using sanger sequencing. Control empty HSV (HSV-e) only expresses GFP. HSV was produced as previously described(Roy et al., 2002). Mice received 35000 pfu/paw (8 µl) intraplantar HSV-e or –iSec1^res^ at days -3 and -1 prior to carrageenan.

Isec1^mut^ was tested for ASO resistance *in vitro* by transfecting Isec1^wt^ or iSec1^res^ into Neuro 2A cells with Lipofectamin 2000, followed by transfection with mis-match (MM) ASO or iSec- targetting ASO (See below: *Antisense oligonucleotide-mediated knockdown*).

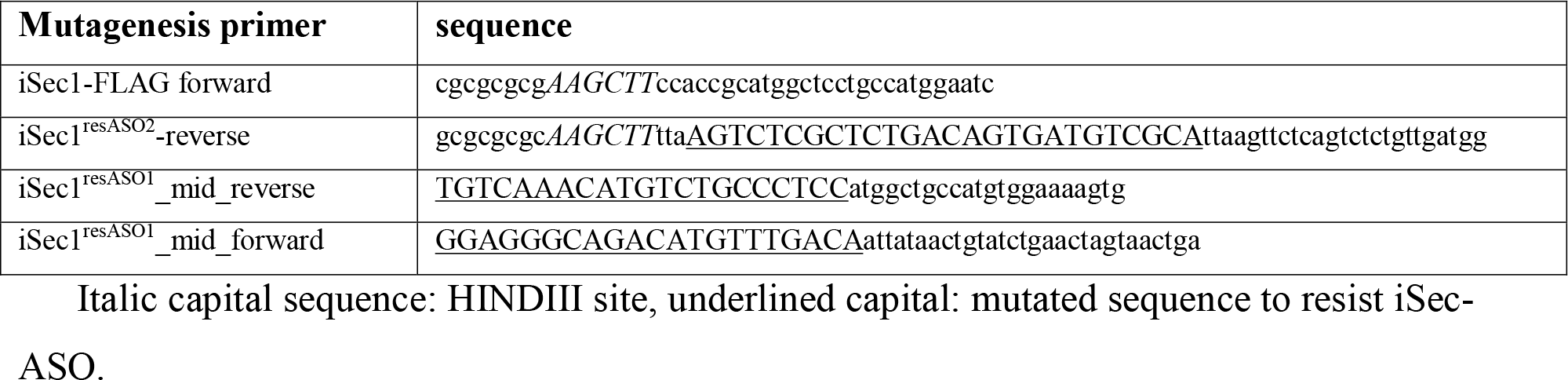

#### Antisense oligonucleotide-mediated knockdown

For in vitro knockdown, we used lipofectamine 2000 (Thermofisher) to transfect N2A- Cd200-iSec1 cells with Mismatch (MM) or iSec1-targetting phosphorothioated antisense oligonucleotides (ASO) according to manufacturer protocol. After 24h, mRNA was isolated with a RNeasy mini kit (Qiagen) and cDNA was generated with iScript (Biorad) according to manufacturer’s protocol. To knockdown iSec1 in sensory neurons in vivo, mice received intrathecal injections of 5 μ iSec-ASO mix (total concentration of 3 μ μ constituting 1:1 mix of iSec-ASO 1 and 2; Sigma-Aldrich) at day 0, 1, 2, 4, and 5. A MM-ASO mix was used as control(Hylden and Wilcox, 1980; Willemen et al., 2018). The following phosphorothioated ASO sequences, that specifically target iSec1/gm609, were used:

| Name | Sequence (5'-3') |
| --- | --- |
| iSec-ASO 1 | [mG]*[mG]*[mU]*[mG]*[mA]*G*G*C*A*G*G*T*T*T*G*[mU]*[mC]*[mC]*[mU]*[mC] |
| iSec-ASO 2 | [mA]*[mC]*[mU]*[mG]*[mG]*C*A*G*T*G*A*T*G*T*C*[mU]*[mC]*[mA]*[mU]*[mU] |
| MM-ASO 1 | [mG]*[mU]*[mG]*[mC]*[mG]*A*T*G*T*A*T*G*C*C*G*[mG]*[mU]*[mG]*[mC]*[mU] |
| MM-ASO 2 | [mG]*[mU]*[mA]*[mU]*[mU]*G*T*T*A*A*C*G*C*G*T*[mA]*[mU]*[mG]*[mC]*[mC] |

[mA] [mG] [mC] [mU] = 2'OMethyl nucleotides to prevent breakdown/toxicity

#### Flow cytometry analysis and cell sorting

DRGs (L3–L5) were collected to analyze infiltrating immune cells. In brief, tissues were gently minced and digested at 37L C for 30L minutes with an enzyme cocktail (5 mg collagenase type I with 2.5 mg trypsin, Sigma Aldrich) in 5 ml DMEM. Cells were stained with various combinations of fluorochrome-labeled antibodies (see subheading ‘antibodies’).

Blood was collected in EDTA tubes (Greiner Bio-One) following heart puncture and erythrocytes were lysed (RBC lysis buffer, eBioscience) before FACS staining.

For vesicles (see isolation of extracellular vesicles), pellets were pooled and stained for CD45, CD11b and CD200 Receptor 1 (CD200R), and MTDR (see subheading ‘antibodies’). Before samples were acquired by LSRFortessa flow cytometer (BD Biosciences) and analyzed with FACSDIVA software, counting beads were added. On average the recovery rate of counting beads (eBioscience) was 44%±3. For all cellular analysis, we used FSC as trigger to identify events. For vesicle analysis, we used FSC or CD45-PB as trigger to identify events.

Advillin^+^ neurons from Advillin^cre-^MitoDendra2^flox^ mice were sorted using an Aria III (BD instruments): Viable^+^Hoechst^+^CD45^-^Advillin^+^, CD45 cells were: iable^+^Hoechst^+^CD45^+^Advillin^-^.

#### Transfer of mitochondria

Bone marrow-derived macrophages were harvested and 2×10^6^ cells were labelled with 20 nM MTDR in 500 µl culture medium. Cells were washed 3 times, counted (NucleoCounter NC-200; Chemometec) and resuspended at a concentration of 120.000 cells/ml medium.

N2As (30.000 cells) were seeded in a 24-well plate and 24h later co-cultured with 12.000 MTDR pre-stained macrophages for 2h and harvested using 1X Trypsin-EDTA (Gibco). Cells were stained for F4/80 and CD11b (see subheading ‘antibodies’). MTDR signal in N2A’s was assessed using the ImageStream MkII (Millipore, Burlington, MA) or flow cytometer (4 laser BD Fortessa, 3 laser BD Canto II).

Primary DRG neurons were cultured as described before (see Cell lines and primary cell cultures)(Eijkelkamp et al., 2013) and co-cultured with MitoDsRed-expressing macrophages. After 16h, co-cultures were fixed and imaged with a Zeiss Axio Observer microscope (Zeiss, Oberkochen).

#### Immunofluorescent staining and detection of mitochondrial transfer in vivo

To monitor mitochondrial transfer in vivo, we injected MTDR-labelled macrophages intrathecal 24 hours prior to killing the mice. Lysm^Cre^ MitoDendra2s^top-flox^ were killed at the indicated days after carrageenan injection. The retrograde marker WGA was injected 2 days prior to killing the mice. Mice were killed by cervical dislocation and lumbar spinal cords and DRGs were collected. Tissues were post-fixed in 4% paraformaldehyde (PFA), cryoprotected in sucrose overnight and embedded in optimal cutting temperature (OCT) compound (Sakura, Zoeterwoude, the Netherlands), and frozen at −80°C.

For immunofluorescence, cryosections (10 μ ) of lumbar DRGs or spinal cords, were stained with primary antibodies overnight at 4°C followed by 2 hours incubation with fluorescent-tagged secondary antibodies (see subheading ‘antibodies’). Nuclei were counterstained with or without 4,6-diamidino-2-phenylindole (DAPI). Immunostaining images were captured with a Zeiss Axio Observer or LSM710 microscope (Zeiss, Oberkochen, Germany) using identical exposure times for all slides within one experiment. Fluorescence intensity was analyzed with ImageJ software. The area of MitoDendra2+ area in the NeuroTrace are was measured using a ImageJ macro that randomizes and anonymizes the images (see supplemental information).

#### RNAscope for iSec1

RNAscope in situ hybridization (ish) multiplex version 2 was performed as instructed by Advanced Cell Diagnostics (ACD). Fresh frozen tissue was cut at 10 µM thickness and immediately stored at -80°C until RNAscope was performed. Slides were immediately transferred to cold (4°C) 4% PFA for 15 minutes. The tissues were then dehydrated in serial increasing ethanol solutions and endogenous peroxidase was blocked for 10 minutes. Samples were incubated with Protease IV for 30 minutes at RT. Samples were incubated with iSec1 probes for 2 hours at 40°C. After probe incubation, the slides were washed in RNAscope wash buffer and returned to the oven for 30 minutes for incubation with AMP-1 reagent. Washes and amplification were repeated using AMP-2 and AMP-3 reagents with a 30-, and 15-minute incubation period, respectively.

#### iDisco, clearing procedure and light sheet imaging

DRGs from adult mice were cleared using iDISCO protocol as described before(Renier et al., 2014). Briefly, animals were perfused with 4% PFA, lumbar dorsal root ganglia were dissected and samples were dehydrated in increasing concentrations (20%, 40%, 60%, 80%, 100%) of methanol solutions. Samples were bleached and rehydrated in decreasing concentrations of methanol solutions. After blocking for 48 hours, samples were incubated with the primary antibodies for 48 hours followed by incubation with secondary antibody for another 48 hours (see subheading ‘antibodies’). After samples were embedded in agarose, they were dehydrated in increasing concentrations of methanol solutions. Samples were incubated overnight in 1 volume of 100% methanol/2 volumes 100% dichloromethane (DCM) anhydrous, washed with 100% DCM and incubated in 100% dibenzyl ether (DBE) for at least one day before imaging. Samples were imaged with an Ultramicroscope II (LaVision BioTec) lightsheet microscope equipped with Imspector (version 5.0285.0) software (LaVision BioTec). The microscope consists of an Olympus MVX-10 Zoom Body (0.63-6.3x) equipped with an Olympus MVPLAPO 2x Objective lens, which includes, dipping cap correction optics (LV OM DCC20) with a working distance of 5.7mm. Images where taken with a Neo sCMOS camera (Andor) (2560×2160 pixels. Pixel size: 6.5 x 6.5 µm^2^). Samples were scanned with a sheet NA of 0.148348 (results in a 5 µm thick sheet) and a step-size of 2.5 µm using the horizontal focusing light sheet scanning method with the optimal number of steps and using the contrast blending algorithm. The following laser filter combinations were used: Coherent OBIS 561-100 LS Laser with 615/40 filter and Coherent OBIS 647-120 LX with 676/29 filter.

#### Real-time RT-PCR

Total RNA was isolated from freshly isolated DRGs (L3-L5) or hind paws using TRizol and RNeasy mini kit (Qiagen, Hilden, Germany). cDNA was synthesized using iScript reverse transcription supermix, according to manufacture protocol (Bio-Rad, Hercules, CA). Quantitative real-time PCR reactions were performed using an I-cycler iQ5 (Bio-Rad, Hercules, CA) as described(Peters et al., 2013) or on a QuantStudio 12K Flex or a StepOnePlus Realtime PCR system (AB Instruments) with SYBR Select Master Mix (Life Technologies). We used 1-5 ng cDNA input per qPCR reaction.

mRNA expression is represented as relative expression = 2^(Ct(average of reference genes)- Ct(target)). For N2As we used the average Ct values of Gapdh and B2M as reference, for mRNA expression in DRG we used the average of TBP and Rictor as reference, for paws *Gapdh* and *B2m*. #1 primers were used for silencing validation in vitro, #2 primers were used for ex vivo mRNA.

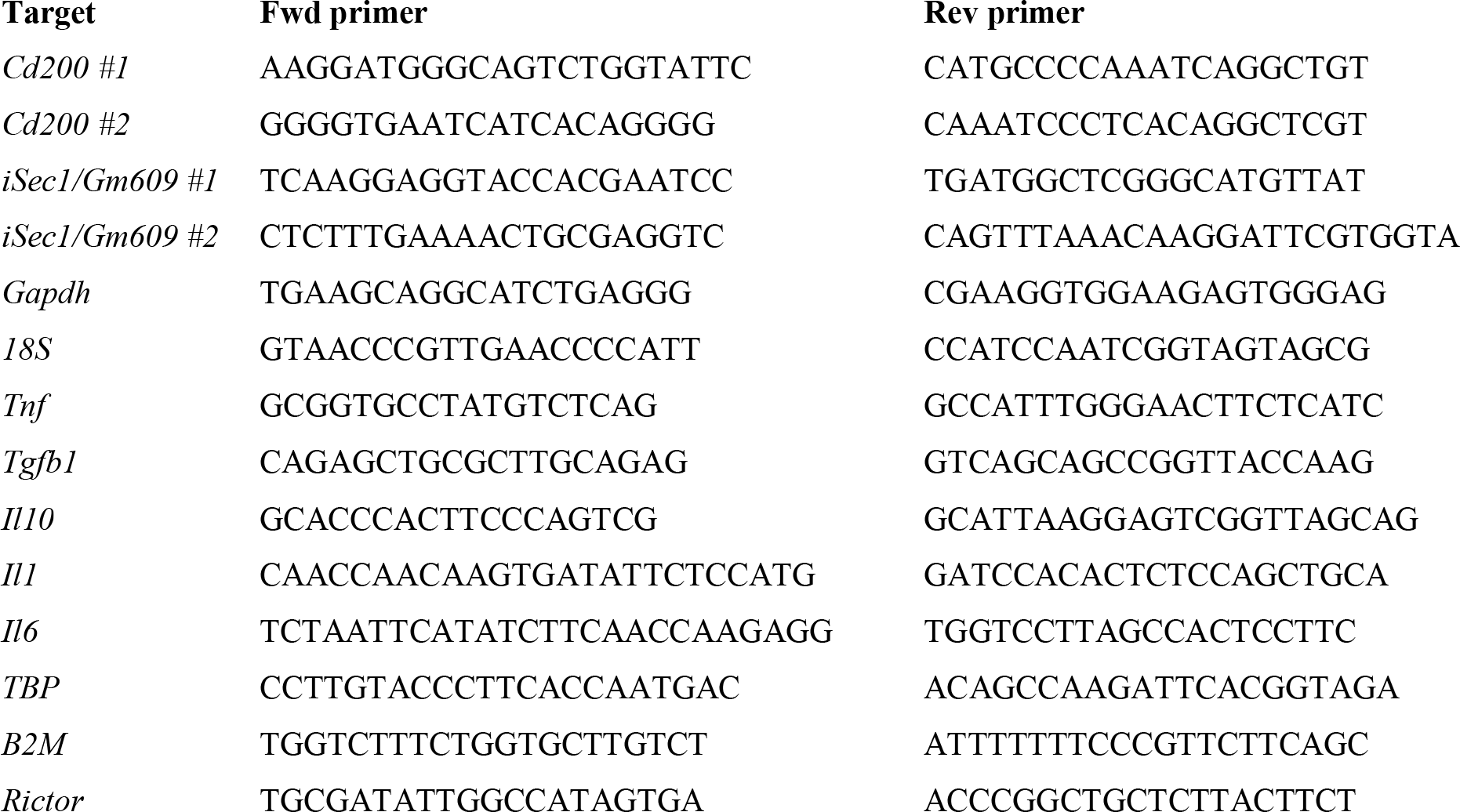

#### Measurement of mitochondrial respiration by Seahorse

Macrophages of WT and *Cd200r*^-/-^ mice were seeded on non-coated XF24 (125k cells) or XF96 (50k cells) plates (Seahorse Bioscience), and grown overnight at 37°C. Next day, cells were washed and placed in Seahorse XF-assay base media (pH 7.4) supplemented with 10 mM glucose and 1 mM sodium pyruvate at 37°C to degas. The Seahorse Bioscience XFe24 Analyzer (Seahorse Bioscience) was used to measure oxygen consumption rates (OCR) under basal conditions, and after sequential addition of oligomycin (1 μ , FCCP (0.2 μ , and rotenone (0.5 μ ), which were injected after cycle 4, 8, and 12, respectively. Each assay cycle consisted of 1.5 minute of mixing, 2 minutes waiting, and 3 minutes of OCR measurements. For each condition, three cycles were used to determine the average OCR under given condition. The measured OCR was normalized for protein content. Five independent experiments were run, each consisting of 3 or more replicates.

DRG OCR analysis as described before (Maj et al., 2017). In brief, primary DRG neurons were cultured as described before (Eijkelkamp et al., 2013) and seeded on poly-d-ornithine/laminin coated XF24 wells plate (15K) and grown overnight at 37°C. Next day, cells were washed and placed in Seahorse XF-assay base media (pH 7.4) supplemented with 4 mM Glutamine, 25 mM glucose and 1 mM sodium pyruvate. OCR was measured under basal conditions.

Mitochondria from WT and *Cd200r^-/-^* macrophages were isolated according to Iuso et al.(Iuso et al., 2017). To measure complex I and complex II driven respiration, 15 μg and 5 μg mitochondria were added in a non-coated XF24 plates, respectively. To measure complex II driven respiration, MAS buffer (220 mM d-Mannitol, 70 mM sucrose, 10 mM KH2PO4, 5 mM MgCl2, 2 mM HEPES, 1 mM EGTA, and 0.2% (w/v) of fatty acid-free BSA, pH 7.2) was supplemented with 10 mM succinate and 2 μM rotenone. For complex I specific respiration, MAS buffer was supplemented with 5 mM malate and 10 mM glutamate. OCR levels were measured under basal conditions, and after sequential addition of ADP (2 mM), oligomycin (3,2 μM), FCCP (4 μM), and antimycin A (4 μM). Each assay cycle consisted of 1 minute of mixing and 3 minutes of OCR measurements. For each condition, three cycles were used to determine the average OCR under given condition.

#### Statistical analysis

All data are presented as mean ± SEM and were analyzed with GraphPad Prism version 8.3 using unpaired two-tailed t tests, one-way or two-way ANOVA, or as appropriate two-way repeated measures ANOVA, followed by post-hoc analysis. The used post-hoc analyses are indicated in each figure. A p value less than 0.05 was considered statistically significant, and each significance is indicated with * or °: p < .05; ** or °° :p < .01; *** or °°°: p < .001; **** or °°°°: p <.0001. All results from statistical analysis are available in the supplemental.

#### Antibodies

| **Target** | **Clone** | **Fluorophore** | **Vendor** | **Catalogue #** |
| --- | --- | --- | --- | --- |
| *Flow cytometry and ImageStream* |  |  |  |  |
| CD115 | AFS98 | APC | eBioscience | 1277550 |
| CD115 | AFS98 | PE-eF610 | eBioscience | 61-1152-80 |
| CD11b | M1/70 | PerCP-Cy5.5 | BioLegend | 101227(8) |
| CD11b | M1/70 | PE | BD Bioscience | 553311 |
| CD11c | N418 | BV785 | BioLegend | 117336 |
| CD19 | 6D5 | PE | BioLegend | 115508 |
| CD200 | OX90 | PE | eBioscience | 12-5200-82 |
| CD200R | OX110 | AF488 | Bio-Rad | MCA2281A488 |
| CD206 | C068C2 | BV650 | eBioscience | 1308615 |
| CD3 | 17A2 | APC | BioLegend | 100236 |
| CD45 | 30-F11 | BV711 | BioLegend | 103147 |
| CD45 | 30-F11 | PB | BioLegend | 103126 |
| CD45 | 30-F11 | APC-eF780 | eBioscience | 47-0451-82 |
| F4/80 | BM8 | FITC | BioLegend | 123108 |
| F4/80 | BM8 | BV510 | BioLegend | 123135 |
| iNOS | CXNFT | APC | eBioscience | 17-5920-80 |
| Ly6C | AL-21 | BV421 | BD Bioscience | 562727 |
| Ly6G | 1A8 | BV785 | BioLegend | 127645 |
| MHCII | M5/114.15.2 | PerCP | BioLegend | 107624 |
| FLAG | L5 | APC | BioLegend | 637308 |
| *Immune fluorescent microscopy and iDISCO* |  |  |  |  |
| F4/80 | C1:A3-1 (rat) | None | Cedarlane | CL8940AP |
| CD206 | Polyclonal goat | None | R&D | AF2535 |
| Iba1 | Polyclonal rabbit | None | Wako | 019-19741 |
| β3-Tubulin | Polyclonal rabbit | None | Abcam | ab18207 |
| Neurofilament M | Polyclonal rabbit | None | Biolegend | 841001 |
| Anti-Goat | Donkey | AF488 | LifeTech | A11055 |
| Anti-Rabbit | Donkey | AF594 | LifeTech | A21207 |
| Anti-Rabbit | Donkey | AF568 | LifeTech | A10042 |
| Anti-Rat | Goat | AF647 | LifeTech | A21247 |

