## Supplemental for "Macrophages transfer mitochondria to sensory neurons to resolve inflammatory pain"

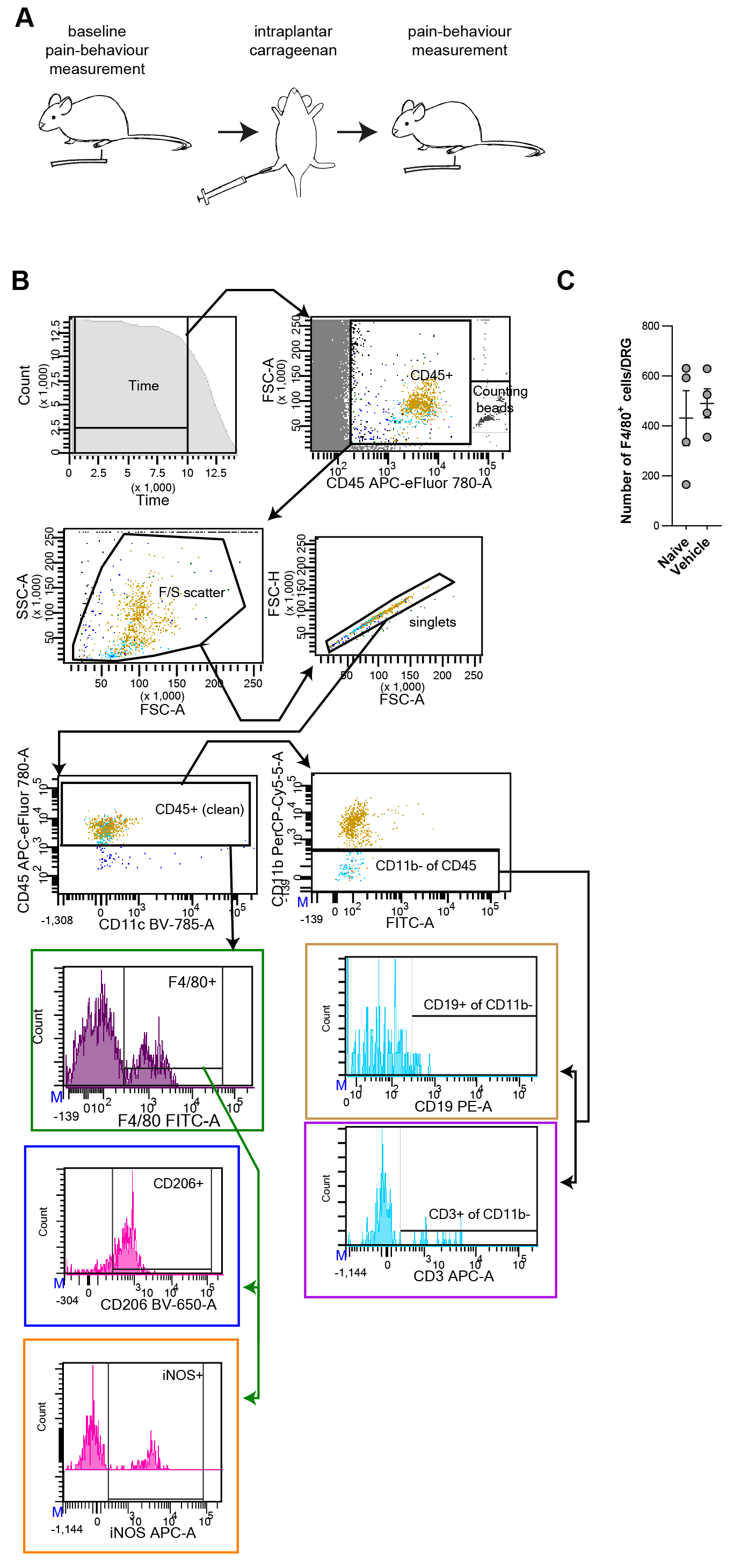

Fig. S1. Carrageenan-induced transient inflammatory pain

(A)	Schematic representation of the carrageenan-induced transient inflammatory pain model. Baseline thresholds were determined three times the first week prior to carrageenan injection and averaged. After intraplantar (i.pl.) carrageenan injection, we measured pain-associated behaviors and signs.

(B)	Gating strategy for monocyte/macrophage phenotyping and T- and B-cells in the DRG. A time gate unified the acquisition analysis, followed by a rough separation of events based on CD45 expression but excluding the counting beads. Cells were further gated by FCS/SSC, single cells and CD45 expression before analysis of CD11b, and F4/80. F4/80+ cells were further assessed for expression of iNOS and CD206; or excluding CD11b+ cells and assessing CD3 and CD19 expression on the CD11b negative cells.

(C)	F4/80 positive macrophage in DRG of control animals or the contralateral lumbar DRG at day 3 after i.pl. injection of carrageenan. Unpaired t-test (ns).

All statistical information can be found in the supplemental data files.

 
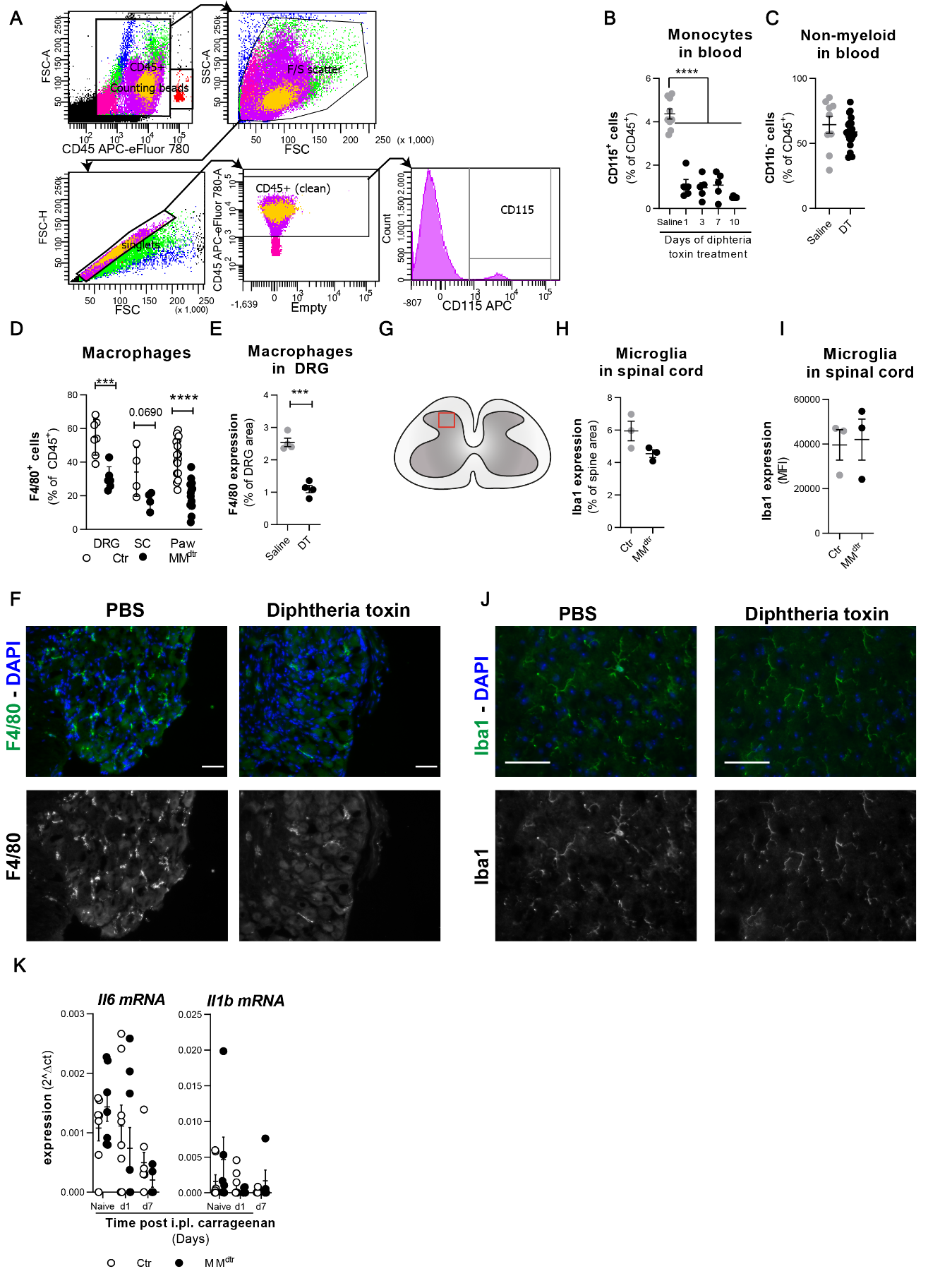

Fig. S2. Depletion controls in MM^dtr^ mice

(A)	Gating strategy for depletion analysis in blood.

(B, C)	Flow cytometry data to validate monocyte depletion in MM^dtr^ mice treated with Diptheria toxin (DT). (B) Percentage of CD115^+^ monocytes in blood to confirm depletion of monocytes, and (C) percentage of non-myeloid cells in blood (CD45^+^CD11b^-^ cells) to confirm depletion specificity. (B) one-way ANOVA, Dunnett; (C) Unpaired t-test with Welch’s correction.

(D)	Percentage of F4/80^+^ macrophages at day 3 in the DRG, spinal cord (SP) and paw, to confirm partial depletion of tissue resident macrophages, for gating see Fig. S1B. Unpaired t-test.

(E, F) Immunofluorescence microscopy analysis of depletion specificity of macrophages versus microglia cells in MM^dtr^ mice after DT treatment. (E) Quantification and (F) example images of F4/80 expression in DRG of MM^dtr^ mice treated with saline or DT. Blue: nuclei, Green: F4/80. Scale bar: 50µM. Unpaired t-test.

(G)	Red square in schematic drawing of lumbar spinal cord indicates the area of dorsal horn analyzed for subfigures H, I and J.

(H-J) Spinal microglia are not depleted after DT treatment. (H, I) Quantification and (J) example images of Iba1^+^ microglia in the spinal cord of MM^dtr^ mice treated with saline or DT. Blue: nuclei, Green: Iba1. Scale bar: 50µM. Significance tested: Unpaired t-test.

(K)	Time course of Il6 and Il1b mRNA expression in Ctr and MM^dtr^ mice treated with carrageenan.

All statistical information can be found in the supplemental data files.

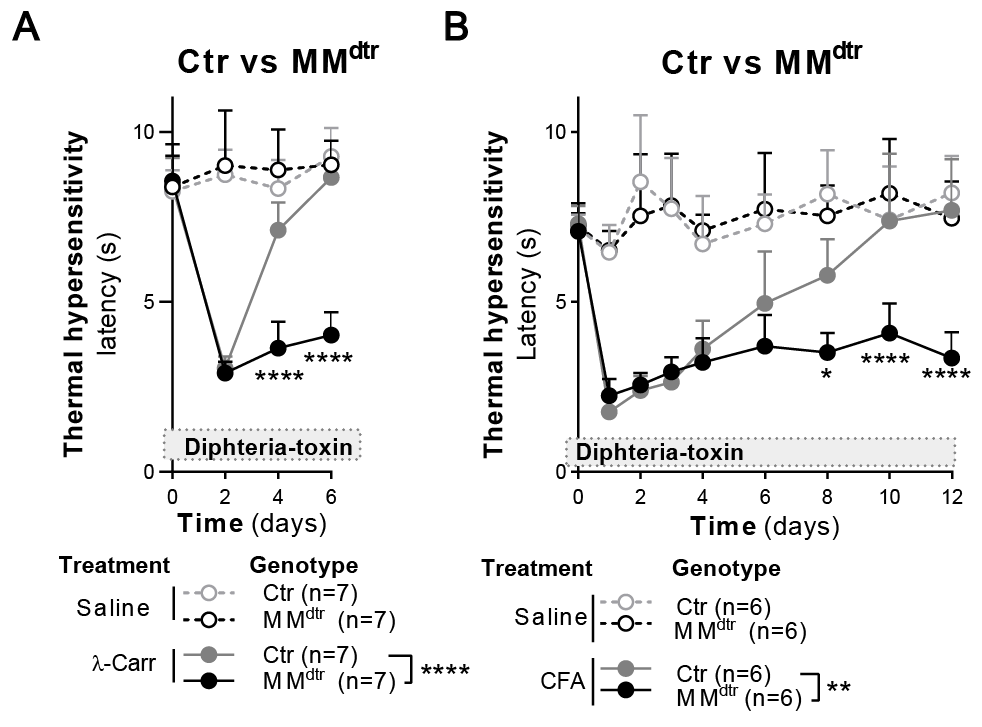

Fig. S3. Course of pain hypersensitivity in MM^dtr^ mice

(A) Course of thermal hyperalgesia in Ctr and MM^dtr^ mice injected with 1% carrageenan in one saline in the other hind paw. 2-way repeated measures ANOVA, Sidak post-hoc comparing carrageenan conditions.

(B) Course of unilateral CFA-induced thermal hyperalgesia in Ctr and MM^dtr^ mice. 2-way repeated measures ANOVA, Sidak post-hoc comparing CFA conditions.

All statistical information can be found in the supplemental data files.

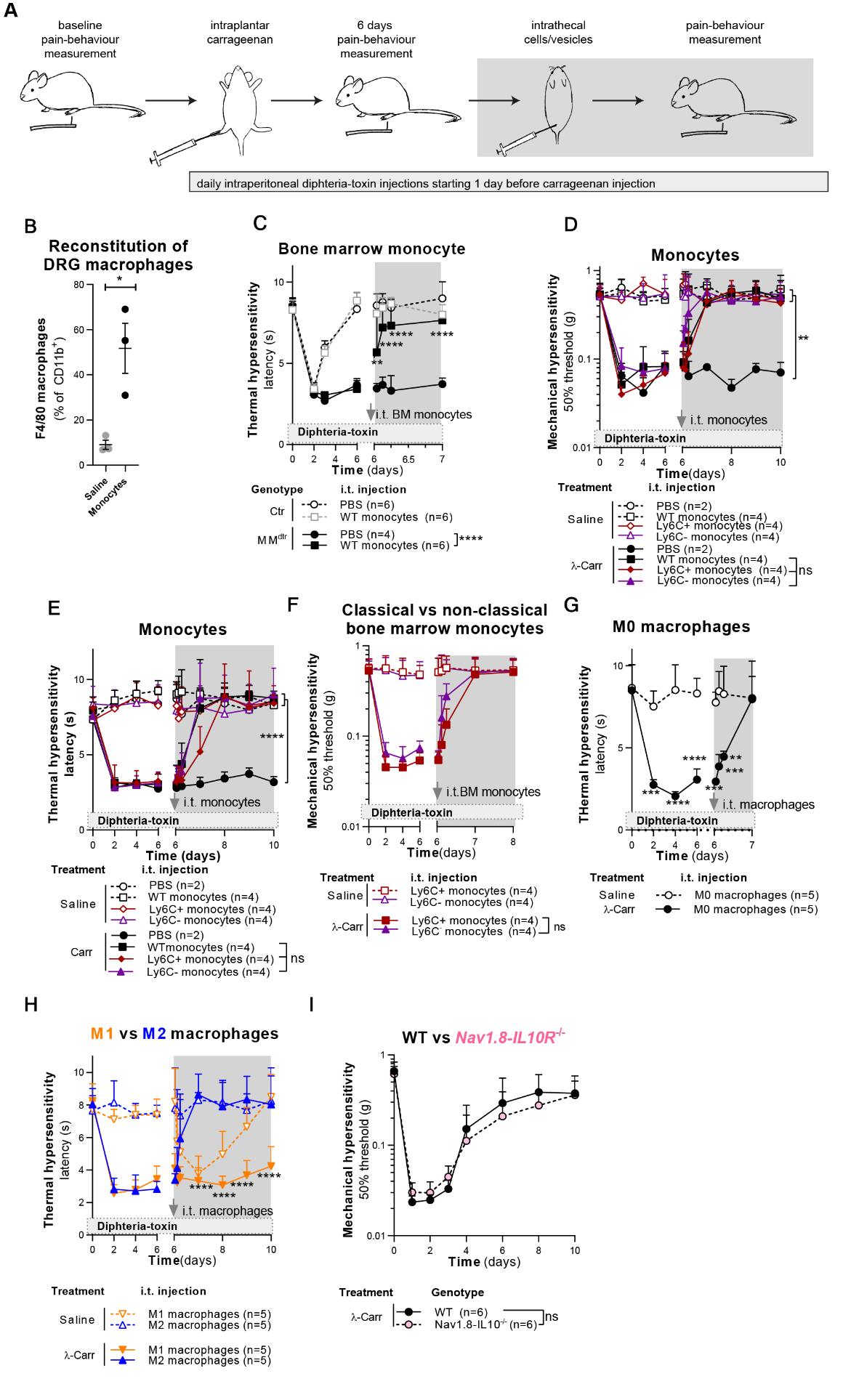

Fig. S4. Depletion of monocytes/macrophages prevents resolution of inflammatory pain

(A)	Schematic representation of the depletion, intraplantar (i.pl.) and intrathecal (i.t.) injections in the pain models.

(B)	Flow cytometry analysis of lumbar DRG (L3-L5) after i.t injection of WT monocytes in MM^dtr^ mice. Gating strategy is indicated in Fig. S2. Unpaired T test.

(C)	Course of carrageenan-induced thermal hyperalgesia in MM^dtr^ mice injected i.t. with CD115^+^ bone marrow monocytes at day 6. 2-way repeated measures ANOVA, Sidak post-hoc comparing MM^dtr^ conditions.

(D, E)	Course of carrageenan-induced mechanical (D) and thermal(E) hyperalgesia in MM^dtr^ mice injected i.t. with splenic CD115^+^ monocytes, ‘classical’ Ly6C^+^ or ‘non-classical’ Ly6C^-^ monocytes or PBS at day 6. 2-way repeated measures ANOVA, Sidak post-hoc comparing carrageenan conditions.

(F)	Course of carrageenan-induced mechanical hyperalgesia in MM^dtr^ mice injected i.t. with ‘classical’ Ly6C^+^ or ‘non-classical’ Ly6C^-^ bone marrow monocytes. 2-way repeated measures ANOVA, Sidak post-hoc comparing carrageenan conditions.

(G)	Course of thermal hyperalgesia in MM^dtr^ mice after i.pl. injection of 1% carrageenan in the left hind paw and i.t. injection of M0 macrophages. 2-way repeated measures ANOVA, with Sidak post-hoc.

(H)	Course of carrageenan-induced thermal hyperalgesia in MM^dtr^ mice injected i.t. with LPS/IFNγ-treated ‘M1’, IL4-treated ‘M2’ macrophages or PBS. 2-way repeated measures ANOVA, Sidak post-hoc comparing carrageenan conditions.

(I)	Course of carrageenan-induced mechanical hypersensitivity in Nav1.8^cre^-*Il10r*^-/-^ litter mates.
2-way repeated measures ANOVA, Sidak post-hoc.

All statistical information can be found in the supplemental data files.

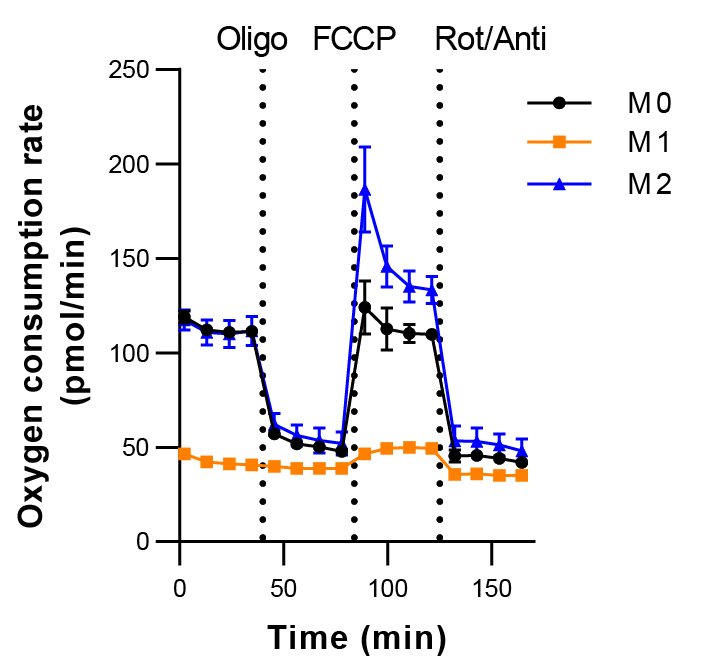

Fig. S5. Macrophage oxidative phosphorylation

Oxygen consumption rate of macrophages (M0), macrophages differentiated with LPS and IFNγ (M1), or with IL4 (M2), as was described before (Van den Bossche et al.)

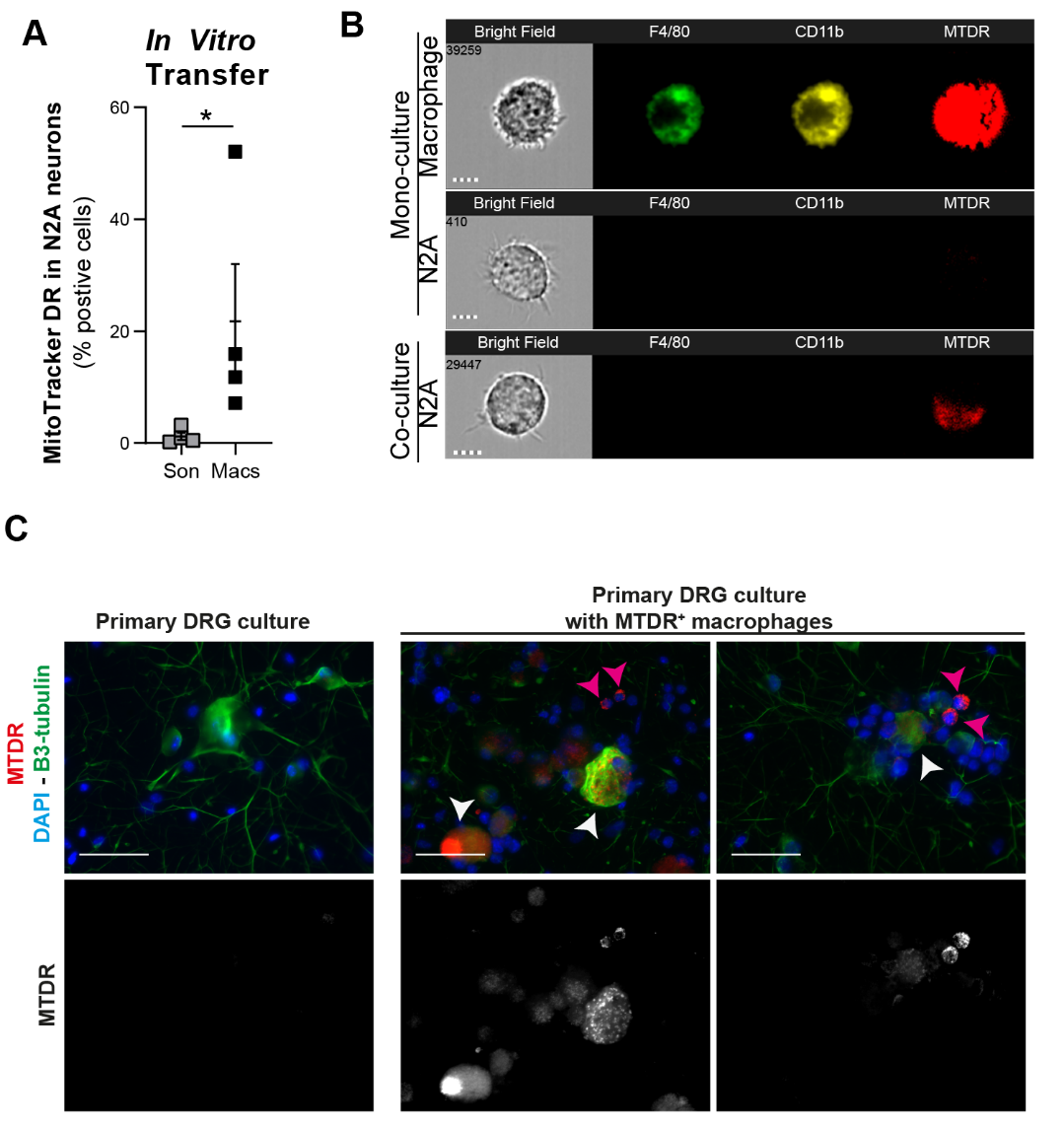

Fig. S6. Macrophages transfer mitochondria to neurons *in vitro*

(A)	Analysis of MitoTracker Deep Red (MTDR) signal in N2A-neurons after co-culture of MTDR-labelled macrophages (macs), or sonicated MTDR-labelled macrophages (Son.) with MTDR-negative N2A-neurons. Co-cultures were stained with CD11b and CD45 to identify macrophages and N2A cells and analyzed by flow cytometry. Mann Whitney test.

(B)	Macrophages labeled with MTDR were co-cultured with N2A for 2 hours and analyzed by image stream (n=2). Co-cultures were stained for CD11b and F4/80 to identify macrophages, and neurons were identified by negative selection. Scale bar: 7µm.

(C)	Macrophages transduced with Mito-dsRed were co-cultured with primary mouse sensory neuron cultures for 16h. Co-cultures were analyzed with immune fluorescence for macrophage-derived mitochondria (red), neurons (β3 tubulin, green), and nuclei (DAPI, blue). Macrophages are indicated with pink arrow heads. Neurons positive for Mito-dsRed are indicated by white arrow heads. n=1. Scale bar: 50µM.

All statistical information can be found in the supplemental data files.

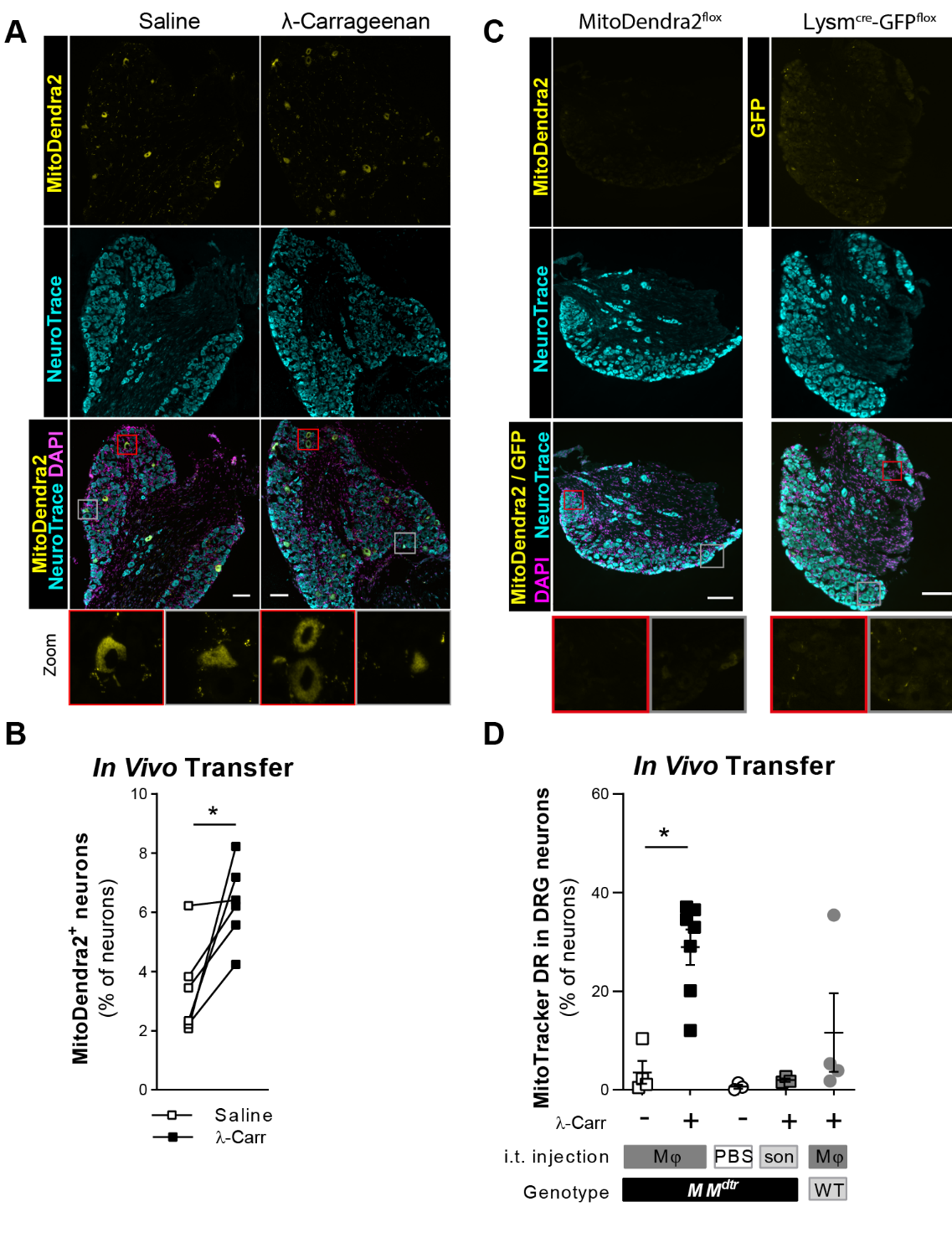

Fig. S7. Macrophages transfer mitochondria to neurons

(A, B)	(A) Example images and (B) quantification of percentage of MitoDendra2+ neurons in the contra- or ipsilateral DRG of LysMcre-MitoDendra2flox mice three days after carrageenan injection. scale bar: 150 µm. n=6. Paired-t Test.

(C)	Example image of MitoDenra2^+^ presence in naïve MitoDendra2^flox^ mouse, or GFP^+^ neurons in *Lysm*^cre^-*GFP*^flox^ mouse.

(D)	Analysis of MTDR signal in sensory neurons in the DRG of MMdtr and Ctr mice. At day 6 after 1% carrageenan (ongoing pain in MM^dtr^, resolved pain in Ctr mice) we i.t. injected PBS, MTDR-labelled macrophages (Mφ), or sonicated MTDR-labelled macrophages (son). After 18h, lumbar DRG were isolated for immunofluorescence analysis and counter-stained with β3-tubulin (cyan, neurons) and DAPI (magenta, nuclei). White arrowheads indicate MTDR^+^ (yellow) neurons. Scale bar: 50µm. Kruskal-Wallis with Dunn post-hoc.

All statistical information can be found in the supplemental data files.

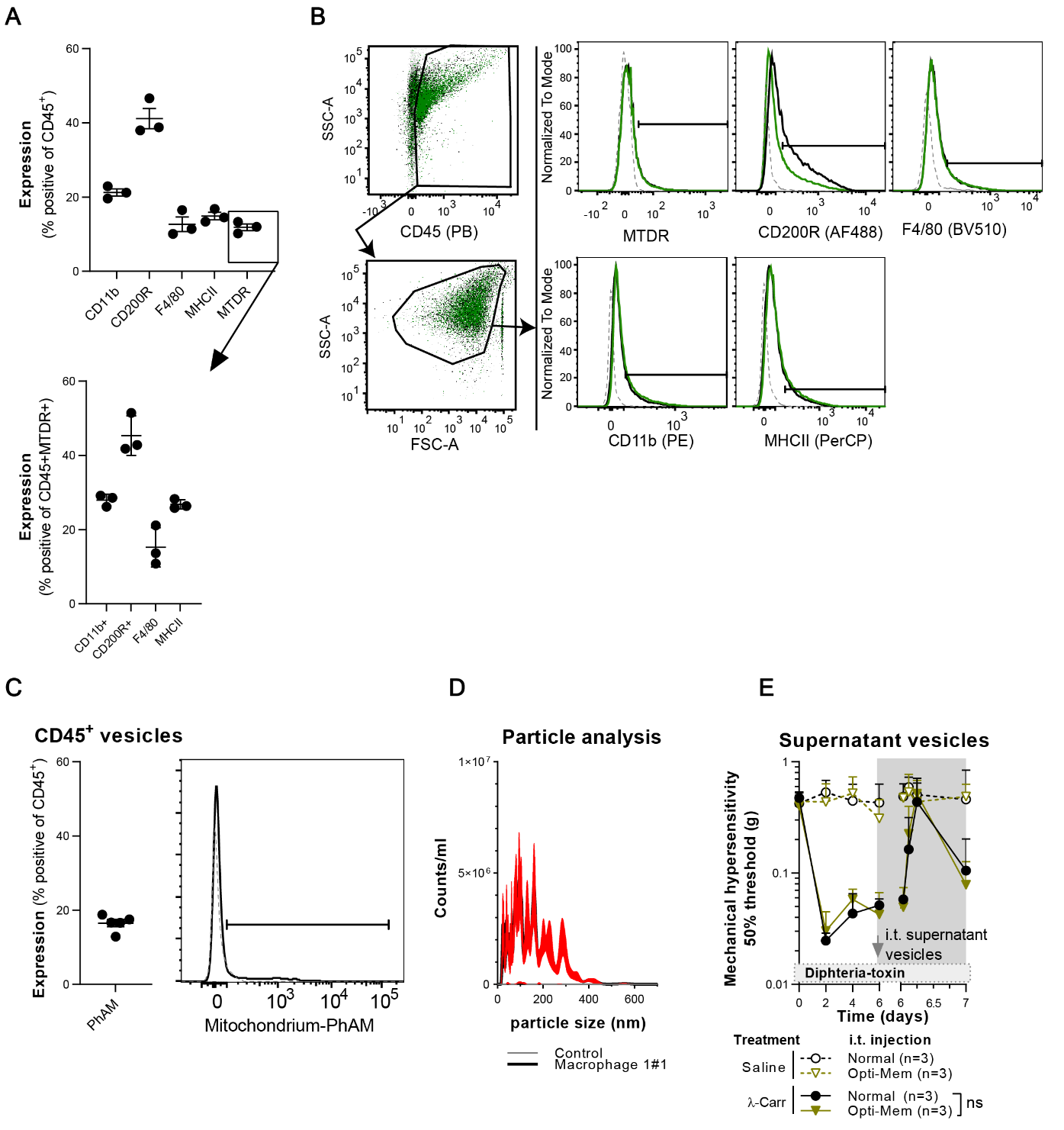

**Fig. S8. Characterization of macrophage-derived extra cellular vesicles.**

(A, B)	Expression of CD11b, F4/80, CD200R and MHCII on CD45^+^ and below CD45^+^MTDR^+^ vesicles. (B) Gating strategy: Events were captured based on FSC or CD45-PB-signal and gated for CD45 expression, followed by an FFC/SSC gate. Dotted line: unstained, solid black: WT, solid green: *Cd200r*^-/-^.

(C)	Expression of Mitochondrion-targeted MitoDendra2 in CD45^+^ vesicles of macrophages, quantified and example histogram.

(D)	Nanoparticle Tracking Analysis (NTA) of supernatant vesicles from WT macrophages. Mean+SEM from 6 technical replicates (red lines are SEM) . Macrophages were cultured in plain Opti-Mem to prevent contamination of FCS particles. As control, medium was ‘cultured’ at 37C in flasks without macrophages.

(E)	Course of mechanical hyperalgesia in MM^dtr^ mice injected with carrageenan in the left hind paw, and saline in the right hind paw. At day 6 mice were injected i.t. with intact macrophage-derived vesicles harvested from macrophage cultured in normal medium (see M&M) or plain Opti-mem. 2-way repeated measures ANOVA, Sidak post-hoc.

All statistical information can be found in the supplemental data files.

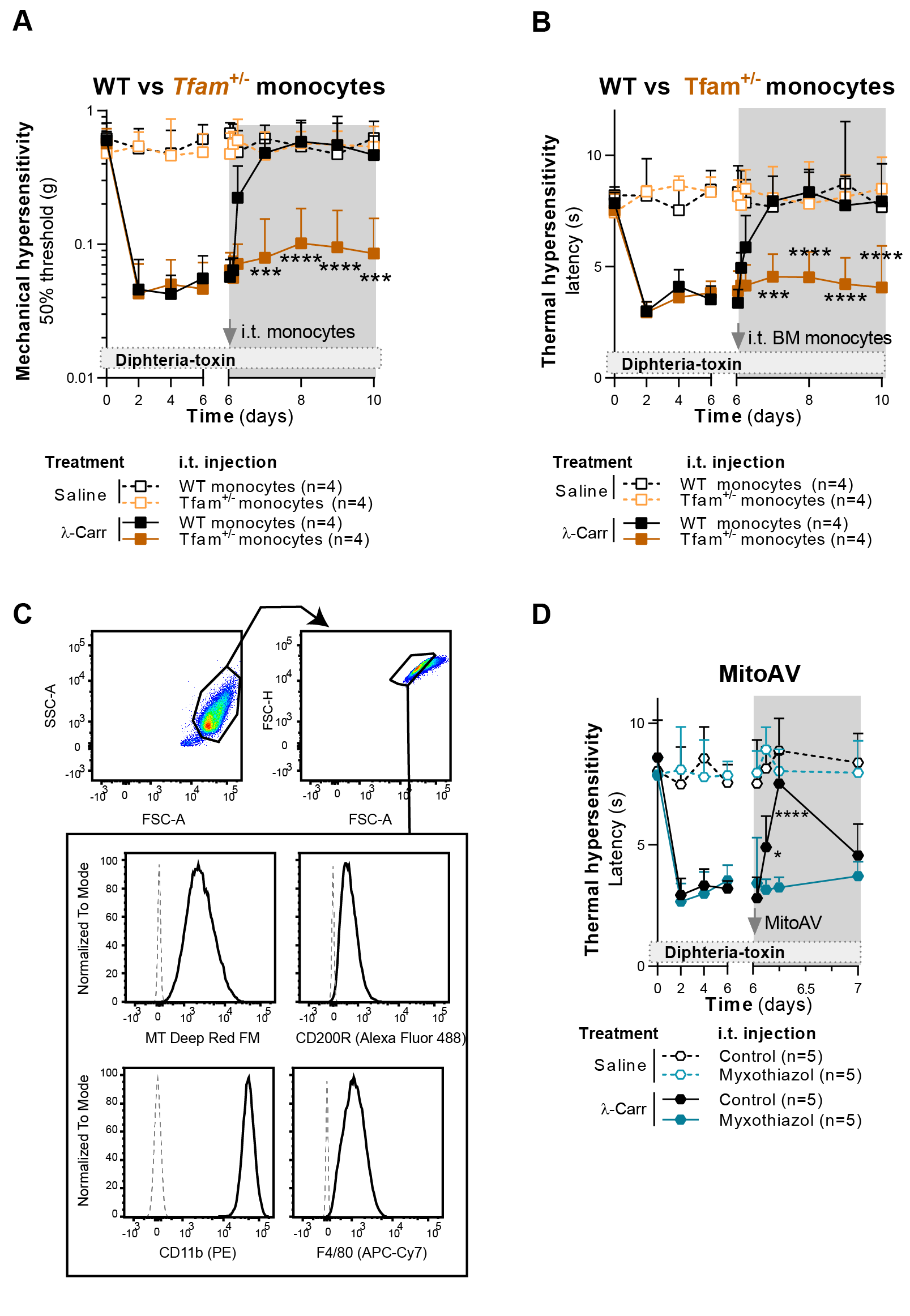

**Fig. S9. Functional mitochondria are required to resolve inflammatory pain**

(A, B)	Course of carrageenan-induced (A) mechanical and (B) thermal hyperalgesia in MM^dtr^ mice injected intra thecal (i.t.) with WT or *Tfam*^+/-^ CD115^+^ monocytes. 2-way repeated measures ANOVA, Sidak post-hoc comparing carrageenan conditions.

(C)	Flow cytometry analysis of isolated mitochondria. Events were gated on FSC/SSC, followed by FCS/FCS-H for singles. Dotted line: unstained, solid black: specific stain.

(D)	Course of carrageenan-induced thermal hyperalgesia in MM^dtr^ mice i.t. injected with functional isolated mitochondria, or mitochondria inhibited with complex III inhibitor myxothizaol. 2-way repeated measures ANOVA, Sidak post-hoc comparing carrageenan conditions.

All statistical information can be found in the supplemental data files.

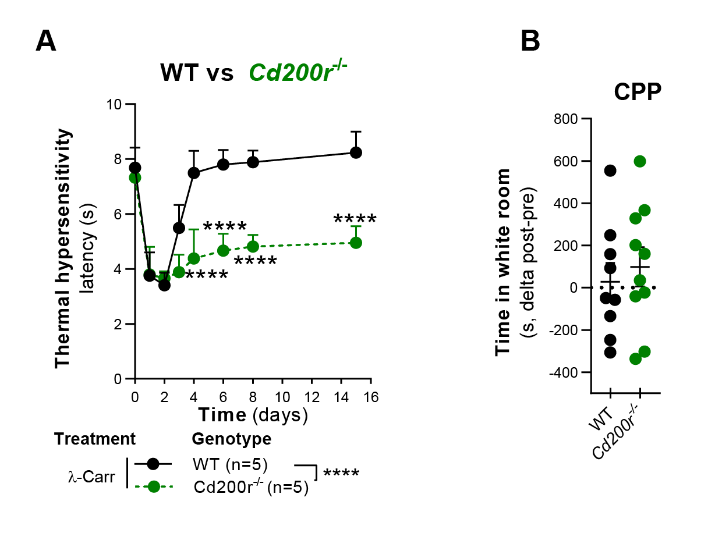

Fig. S10. CD200R is required for the resolution of inflammatory pain

(A)	Course of carrageenan-induced thermal hyperalgesia in WT and *Cd200r*^-/-^ littermates. 2-way repeated measures ANOVA, Sidak post-hoc.

(B)	Gabapentin-induced place preference conditioning at day 16 after unilateral saline injection in the hind paws. Conditioning efficiency is depicted as the difference in time (seconds, s) spent in a white room pre- and post-conditioning. Unpaired t-test.

All statistical information can be found in the supplemental data files.

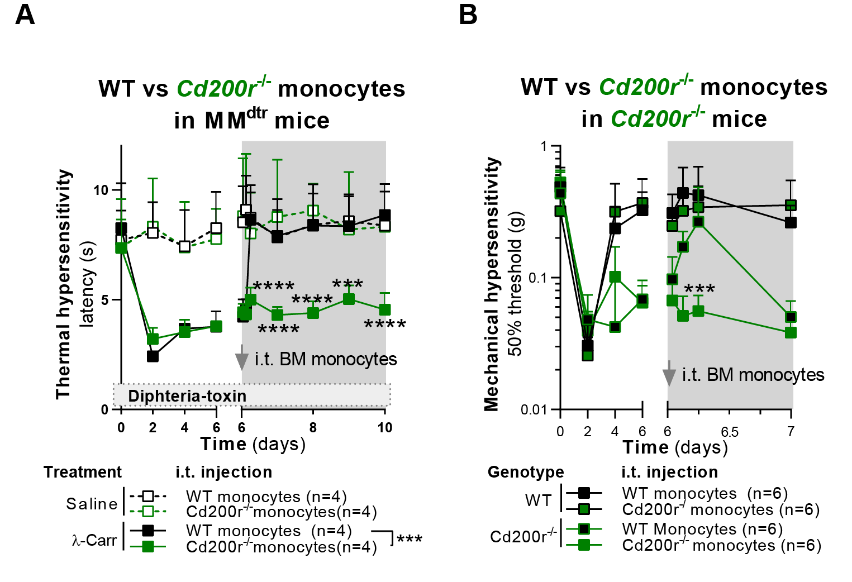

Fig. S11. CD200R is required for the resolution of inflammatory pain

(A)	Course of carrageenan-induced thermal hyperalgesia in MM^dtr^ mice intra thecal (i.t.) injected with WT of *Cd200r^-/-^* CD115^+^ monocytes. 2-way repeated measures ANOVA, Sidak post-hoc comparing carrageenan conditions.

(B)	Course of carrageenan-induced mechanical hyperalgesia in *Cd200r*^-/-^ mice i.t injected with WT of *Cd200r*^-/-^ CD115^+^ monocytes. 2-way repeated measures ANOVA, Sidak post-hoc comparing carrageenan conditions.

All statistical information can be found in the supplemental data files.

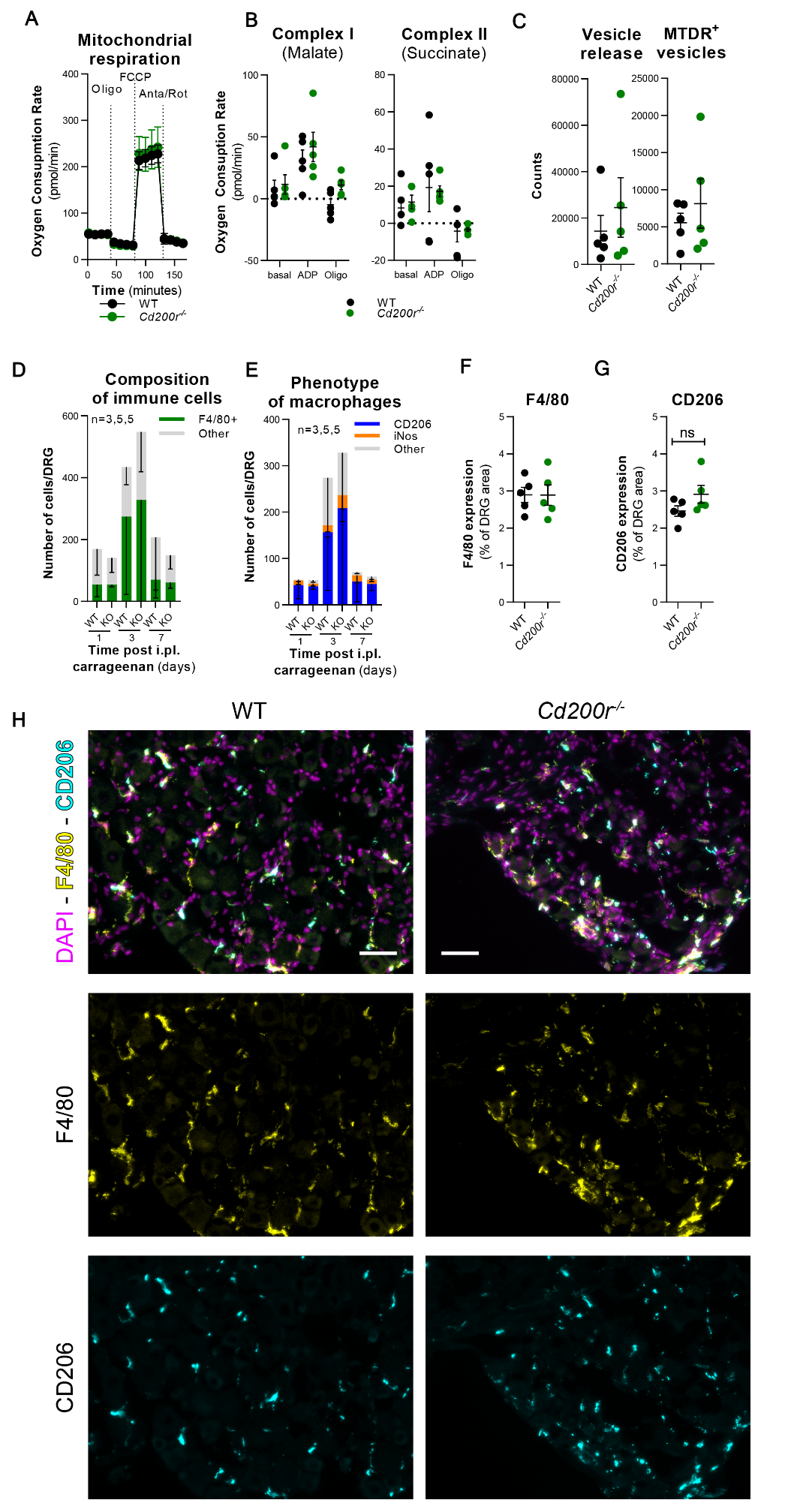

Fig. S12. *Cd200r*^-/-^ have normal vesicles, mitochondrial respiration and macrophage infiltration into DRG

(A)	Mitochondrial respiration in WT or *Cd200r*^-/-^ macrophages. 2-way repeated measures ANOVA, Bonferroni post-hoc.

(B)	Mitochondrial respiration in MitoAV from WT or *Cd200r*^-/-^ macrophages assessed by extracellular flux assay. 2-way repeated measures ANOVA, Bonferroni post-hoc.

(C)	Analysis of vesicle content of WT or *Cd200r*^-/-^ macrophage-conditioned medium. Unpaired t-test.

(D, E)	Flow cytometry analysis of monocytes/macrophages infiltrating the DRG. 2-way repeated measures ANOVA, Bonferroni post-hoc.

(F, G, H) Quantification and example image of F4/80 and CD206 expression in DRG by immune fluorescence in naïve WT and *Cd200r*^-/-^ mice. Scale bars: 50µm. Unpaired t-test.

All statistical information can be found in the supplemental data files.

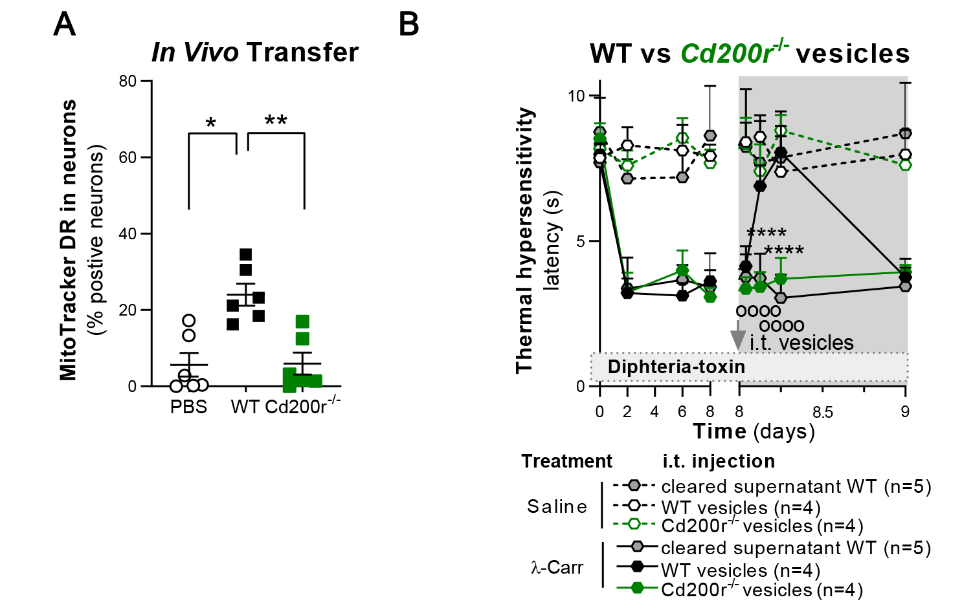

Fig. S13. *Cd200r*^-/-^ macrophages cannot resolve inflammatory pain.

(A) In vivo MTDR transfer from WT or *Cd200r*^-/-^ MTDR-labelled macrophages to DRG neurons of *Cd200r*^-/-^ mice. At day 6 after carrageenan injection, macrophages or PBS were injected i.t. and after 18h DRG were isolated and stained as described for Fig. 2B. Kruskal-Wallis, Dunn post-hoc.

(B) Course of carrageenan-induced thermal hyperalgesia in MMdtr mice i.t injected with vesicles derived from WT of *Cd200r*^-/-^-macrophage-conditioned medium. 2-way repeated measures ANOVA, Dunnet post-hoc comparing carrageenan conditions.

All statistical information can be found in the supplemental data files.

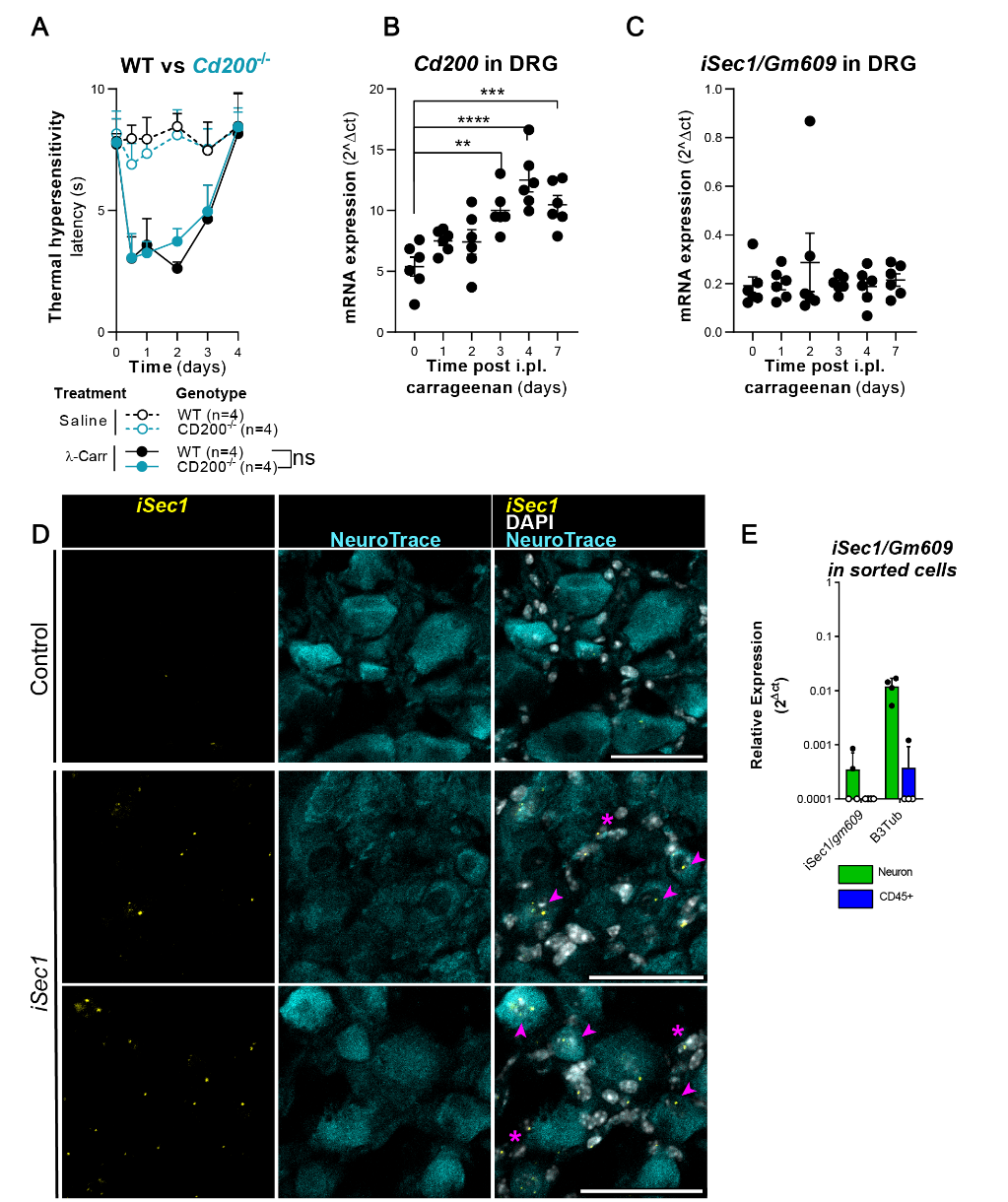

Fig. S14. Characterization of iSec1 expression

(A)	Course of carrageenan-induced thermal hyperalgesia in WT and *Cd200*^-/-^ littermates. 2-way repeated measures ANOVA, Sidak post-hoc comparing carrageenan conditions.

(B)	*Cd200* mRNA expression in DRG of WT mice at indicated days after intra plantar (i.pl) injection of carrageenan. Ordinary ANOVA, Dunnett post-hoc comparing vs time point 0.

(C)	*iSec1/gm690* mRNA expression in DRG of WT mice at indicated days after intra plantar (i.pl) injection of carrageenan. Ordinary ANOVA, Dunnett post-hoc comparing vs time point 0.

(D)	Representative pictures of mRNA expression using in-situ hybridization (RNAScope). No targeting probe (control) and isec1 specific probes (iSec1) were combined with NeuroTrace staining. Arrows represent neurons positive for iSec1, asterisks (*) represent other cells.

(E)	*iSec1/gm609* and *Tubb3* mRNA expression relative to *Gapdh* and *18S* in FACSorted Addvilin^+^ neurons and CD45^+^ immune cells. Open circles (o) represent samples with no amplification.

All statistical information can be found in the supplemental data files.

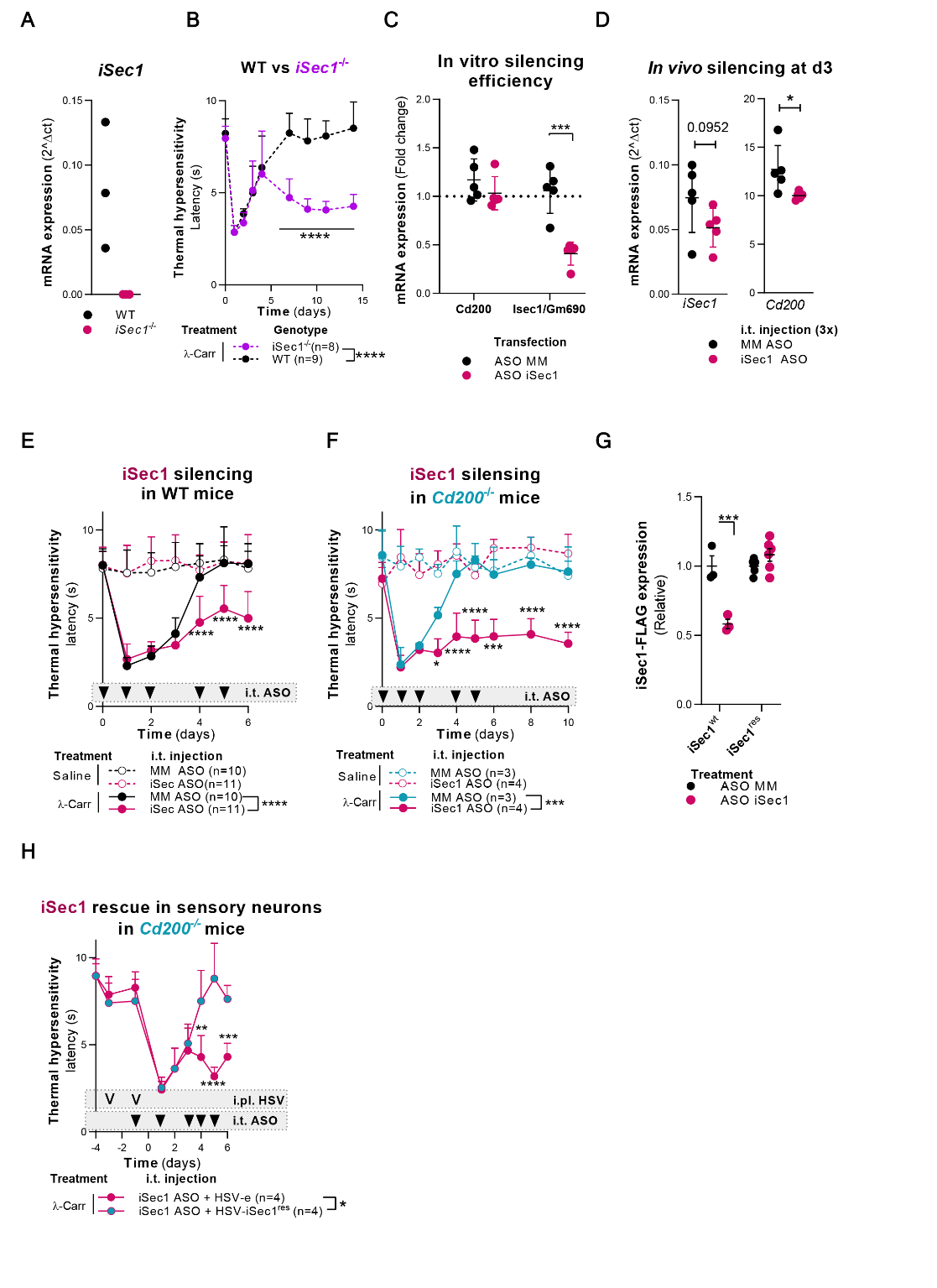

Fig. S15. iSec1 is required for resolution of inflammatory pain

(A)	iSec1 mRNA expression relative to in DRG of WT and *iSec1*^-/-^ mice.

(B)	Course of carrageenan-induced thermal hyperalgesia in WT mice that are control- or iSce1^-/-^ mice. 2-way repeated measures ANOVA, Sidak post-hoc.

(C)	*Cd200* and *iSec1/gm609* mRNA expression after silencing of iSec1/gm609 in neuronal N2A cells ectopically expressing Cd200 and iSec1/gm609. n=5 from 2 experiments. 2-way repeated measures ANOVA, Sidak post-hoc

(D)	iSec1/*gm609* and *Cd200* mRNA expression after silencing of iSec1/*gm609* in DRG of WT mice treated with mismatch (MM-ASO) or iSec1-targeting Antisense Oligo nucleotides (iSec1-ASO). Unpaired t-test.

(E)	Course of carrageenan-induced thermal hyperalgesia in WT mice that are control- or iSec1/*gm609* silenced by ASO. 2-way repeated measures ANOVA, Sidak post-hoc comparing carrageenan conditions.

(F)	Course of carrageenan-induced thermal hyperalgesia in *Cd200*^-/-^mice that are control- or iSec1/*gm609* silenced by ASO. 2-way repeated measures ANOVA, Sidak post-hoc comparing carrageenan conditions.

(G)	Expression of iSec1^wt^-flag (n=3 from 1 experiment) and iSec1^res^-flag (n=6 from 2 experiments) as assessed by flow cytometry. 2-way ANOVA, Sidak post-hoc.

(H)	Course of carrageenan-induced thermal hyperalgesia in *Cd200^-/-^* mice that are iSec1/gm609 silenced by ASO, while i.pl. infected with HSV-e or HSV-iSec1. 2-way repeated measures ANOVA, Sidak post-hoc.

All statistical information can be found in the supplemental data files.

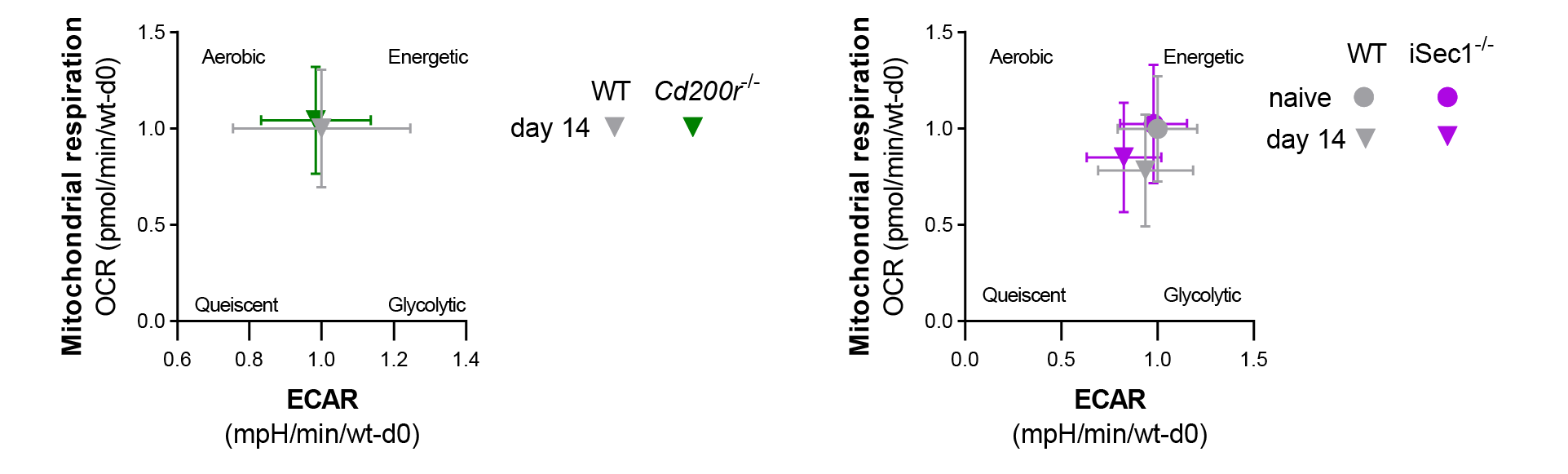

Fig. S16. Energy maps of DRG-cultures from Cd200r^-/-^ and iSec1^-/-^ mice.

Energy maps of DRG cultures from naïve *iSec1^-/-^* and WT mice, and intra plantar carrageenan injected *iSec1*^-/-^, *Cd200r*^-/-^ with appropriate WT controls at day 14. Data are normalized per experiment to OCR/ECAR values on day 0.

Movie S1. 3D render of control DRG

Video render of a lumbar DRG from a saline treated animal. iDISCO immunofluorescence staining for macrophage marker F4/80 (red) or neuronal marker Neurofilament M (green). The effective magnification for all images was 13.6x (zoombody * objective+dipping lens = 6.3x*2.152x).

Movie S1. 3D render of carrageenan DRG

Video render of a lumbar DRG 1 day after 1% carrageenan injection. iDISCO immunofluorescence staining for macrophage marker F4/80 (red) or neuronal marker Neurofilament M (green). The effective magnification for all images was 13.6x (zoombody * objective+dipping lens = 6.3x*2.152x).
